# Longitudinal magnetic resonance imaging and spectroscopy in a mouse model of cuprizone-induced demyelination

**DOI:** 10.64898/2026.02.18.706313

**Authors:** Esther Walters, Davide Di Censo, Elena Samoylenko, Eugene Kim, Sally Loomis, Charalampos Papaonisiforou, Camilla Simmons, Grace Flower, Katarina Ilic, Eilidh MacNicol, Maria Elisa Serrano Navacerrada, Luiza-Simona Damoc, David Virley, Steve Williams, Nicola Hamilton-Whitaker, Andrew McCreary, Diana Cash

**Author notes:** Contributed equally (Esther Walters, Davide Di Censo, Andrew McCreary, Diana Cash).

## Abstract

The cuprizone (CPZ) lesioned mouse is a widely used model of demyelination and remyelination, but most studies rely on histology at terminal timepoints, limiting understanding of disease dynamics. Here, we present a longitudinal multimodal magnetic resonance imaging and spectroscopy (MRI/MRS) study of CPZ-induced pathology, pooling control arms from three independent experiments (n = 40). Mice were imaged at baseline, then exposed to 0.2% cuprizone for 5 weeks and repeatedly imaged at days 24, 35, 49, 63 and 77 after the start of CPZ. Imaging included multiparametric mapping (MPM), diffusion tensor imaging (DTI), tensor-based morphometry (TBM), and single-voxel MRS in the corpus callosum. Histology (MBP, silver, GFAP, Iba1) was performed at selected timepoints for validation. An additional group of 18 CPZ-lesioned mice were imaged ex vivo using a different higher resolution MRI protocol and compared against 19 non-CPZ controls.

MPM-derived MTsat*δ* and R1 reductions indicated robust demyelination in the corpus callosum and deep cerebellar nuclei by day 24, expanding to cortex and hippocampus by day 35. Partial recovery was observed by day 77 but changes persisted, consistent with histological evidence. TBM revealed dynamic volumetric alterations, including hippocampal and cerebellar expansion alongside cortical and subcortical shrinkage, persisting beyond CPZ cessation. DTI demonstrated early (days 24-35) decreases in FA and MD, followed by complex trajectories consistent with microstructural disruption and partial repair. MRS detected early increases in GABA, glutamine, taurine, and glutathione, with corresponding decreases in NAA, while inositol showed a biphasic decrease–increase profile, likely reflecting acute astrocytic dysfunction followed by gliosis – neuroinflammatory processes that were corroborated by immunohistochemistry.

Together, these results demonstrate that multimodal MRI/MRS sensitively captures widespread, dynamic, and only partially reversible pathology in CPZ-treated mice. Longitudinal imaging provides a non-invasive, translational approach to characterising demyelination, gliosis, and remyelination, offering a powerful alternative to histology for preclinical studies and longitudinal therapeutic screening.

## Introduction

The cuprizone lesioned mouse model is a widely used experimental paradigm for studying demyelination and remyelination in the brain (Vega-Riquer et al., 2019; Zhan et al., 2020; Kipp, 2024). Cuprizone is a copper chelator that is administered to mice in powdered rodent chow at 0.2% for 4–6 weeks. Cuprizone administration results in widespread brain pathology, including demyelination, axonal damage, gliosis, and metabolic changes (Gudi et al., 2009, 2014; Castillo-Rodriguez et al., 2022; Zirngibl et al., 2022; Beliard et al., 2025). Following cessation of cuprizone administration the mice exhibit spontaneous remyelination (Leo and Kipp, 2022; Friesen et al., 2024).

Although cuprizone has been used for decades to consistently and reliably induce this pathology in mice (Carlton, 1967), its precise mechanism of action remains unclear (Morgan et al., 2022; Kipp, 2024). Most studies use the standard acute cuprizone protocol described above and have focused exclusively on the corpus callosum, often examining a single timepoint—typically the peak of pathology, at around 5 weeks of cuprizone exposure (Chrzanowski et al., 2019; Kipp, 2023; Friesen et al., 2024; Kipp, 2024). Remyelination is generally thought to begin rapidly – within a week after cuprizone withdrawal and, according to some reports, may be largely complete within 6 weeks (Matsushima and Morell, 2001; Kipp et al., 2009; Sachs et al., 2014; Tagge et al., 2016; Vega-Riquer et al., 2019). However, this may reflect methodological sensitivity as not all studies find remyelination to be complete even after 6 weeks (Crawford et al., 2009; Hertanu et al., 2023).

The main value of the cuprizone model lies in its simplicity and reproducibility, making it highly suitable for compound screening and candidate drug development (Kipp et al., 2009; Praet et al., 2014; Vega-Riquer et al., 2019; Zirngibl et al., 2022; Friesen et al., 2024; Kipp, 2024). Histological methods remain the gold standard for detecting specific cellular and molecular changes; however, they are laborious, require animal sacrifice, and therefore limit longitudinal studies (Zhan et al., 2020; Kipp, 2024). While histology is ideal for mechanistic insight, it is not easily translatable to clinical practice (Wood et al., 2016; Omer et al., 2025).

By contrast, imaging is the clinical method of choice for the diagnosis and monitoring of myelination changes and neuroinflammation associated with multiple sclerosis (MS), the principal disorder studied with this preclinical model (Kaunzner and Gauthier, 2017). Magnetic resonance imaging (MRI) has long been used in MS patients to detect focal lesions and track lesion activity (Calvi et al., 2022). Owing to its non-invasive nature, high sensitivity, and suitability for repeated measures, MRI is an ideal tool for diagnosis, monitoring disease progression, and assessing treatment response (Rocca et al., 2024). In the clinic, T2-weighted and fluid-attenuated inversion recovery (FLAIR) scans are used to detect and quantify focal white matter lesions, thereby providing an overall measure of lesion burden (Neema et al., 2007). Chronic T1 hypointense lesions are interpreted as markers of severe tissue injury (Dupuy et al., 2015) while T1-weighted imaging with gadolinium contrast enables the identification of active lesions, reflecting ongoing inflammation and blood–brain barrier disruption (Rovira et al., 2024).

More recently, advanced MRI has revealed global brain alterations beyond discrete lesions, likely reflecting axonal degeneration, neuronal and synaptic loss, and inflammation-related changes (Hemond and Bakshi, 2018; Nistri et al., 2024). These newer quantitative MRI techniques include magnetisation transfer imaging, diffusion-based methods, and myelin water imaging, and allow for the assessment of myelin integrity and microstructural damage (Mancini et al., 2020; Rahmanzadeh et al., 2022; Khormi et al., 2023). Volumetric analyses of brain atrophy (Lie et al., 2022) provide biomarkers of global and regional neurodegeneration (Rocca et al., 2024). Although these measures are not diagnostic at the individual level, cohort-based analyses provide valuable information about brain health, disease progression, and interactions with normal processes such as ageing (Tokarska et al., 2023).

Magnetic resonance spectroscopy (MRS) has additionally been used in MS and other neurological disorders to quantify MR-visible brain metabolites that may serve as biological markers or diagnostic indicators. To this end, several metabolites have emerged as promising markers in clinical MS (Swanberg et al., 2019). These include N-acetylaspartate (NAA) and N-acetylaspartylglutamate (NAAG), which are both predominantly neuronal and known to serve as acetate reservoirs for myelin lipid synthesis and whose reductions in MS are thought to mark disrupted neuroaxonal integrity. Other metabolites frequently reported to change, though inconsistently in direction, include choline-containing compounds (linked to phospholipid membrane turnover), creatine (cellular energy metabolism), and inositol and lactate (associated with glial and inflammatory processes).

However, despite numerous studies, findings remain heterogeneous and no robust consensus biomarker has emerged. This variability reflects several factors, including voxel placement (MRS is typically acquired from a small brain volume rather than the whole brain), acquisition and analysis methods (hardware, quantification approach, spectral quality, water suppression, T1/T2 effects), and the limited sensitivity of clinical field strengths (*<*3 T) to detect low-concentration metabolites such as GABA and glutamate. A further major source of variability is the clinical heterogeneity of MS itself, with different phenotypes (relapsing-remitting, primary progressive, secondary progressive) and variable lesion distribution across brain and spinal cord. These challenges complicate voxel placement and hinder group-level averaging across patients. By contrast, preclinical MRS in models such as cuprizone partially mitigates these limitations: higher magnetic fields (e.g., 9.4 T) provide greater sensitivity, and cuprizone-induced demyelination is regionally consistent—most prominent in the corpus callosum—making voxel placement more straightforward.

Together, MRI and MRS offer sensitive, non-invasive biomarkers of pathology that can be translated between clinical and preclinical research. Notably, however, despite the broad availability of small-animal scanners and improved analytical pipelines, surprisingly few whole-brain longitudinal MR studies have been performed in the cuprizone model, even though it is one of the best-characterised and most widely used mouse models of demyelination.

Building upon our previous ex vivo characterisation on the CPZ mouse model (Wood et al., 2016), here we conducted several in vivo longitudinal multimodal MR studies in collaboration with academic and industrial colleagues to evaluate the therapeutic efficacy of several compounds. Each study followed a standardised imaging protocol combining multiparametric mapping (MPM), diffusion tensor imaging (DTI), and single-voxel MRS of the corpus callosum, all acquired within a 1.5 h scan session per mouse at each time point. MPM provided MT-, proton density-, and T1-weighted images, from which we calculated MTsat*δ* and *R*_1_ as our best approximation of myelin content (see Figure S2 and (Friesen et al., 2024)). DTI yielded parameters including FA, MD, AD, and RD, while whole-brain regional volumes were assessed using tensor-based morphometry (TBM). Finally, MRS enabled quantification of ten metabolites.

From these studies, we found that multimodal MR is a powerful and efficient method to non-invasively monitor pathology in cuprizone-treated mice. To extend these findings and gain a clearer picture of the underlying biology, we pooled the non-drug control animals from three independent experiments and analysed them collectively. The results of this pooled analysis are presented here, while the therapeutic intervention studies will be described elsewhere.

## Materials and Methods

### Animals

All animals were treated in accordance with the Animals (Scientific Procedures) Act 1986 and King’s College London local ethical guidelines. For the in vivo study, n = 40 C57BL/6JOlaHsd mice (Envigo, UK) aged 8-10 weeks old were used. For the ex vivo study, n = 37 mice on a C57BL/6J background aged 8-10 weeks were used. Sex and sample size breakdowns of study animals are shown in Table 1. Some male animals from the in vivo study and a number of no-cuprizone controls were also used for the post-mortem histological evaluation; a breakdown of which is shown in Table 2.

**Table 1:**
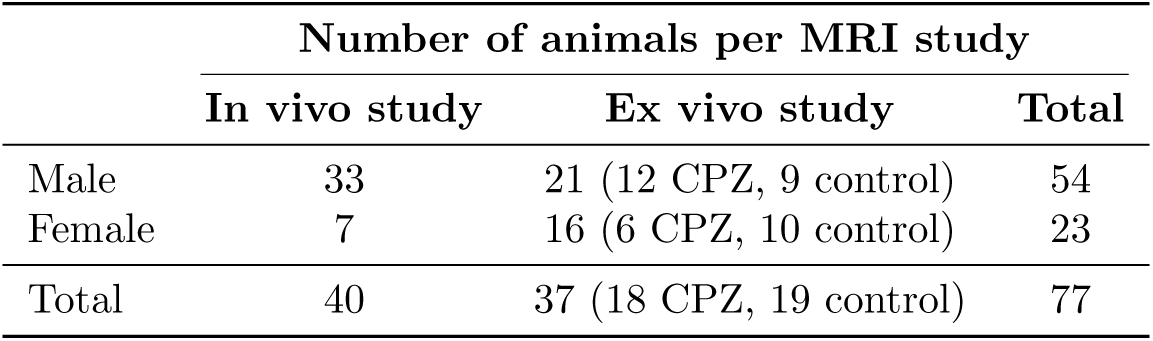
Breakdown of study animals by sex and numbers.

|  | Number of animals per MRI study |  |  |
| --- | --- | --- | --- |
|  | In vivo study | Ex vivo study | Total |
| Male | 33 | 21 (12 CPZ, 9 control) | 54 |
| Female | 7 | 16 (6 CPZ, 10 control) | 23 |
| Total | 40 | 37 (18 CPZ, 19 control) | 77 |

**Table 2:**
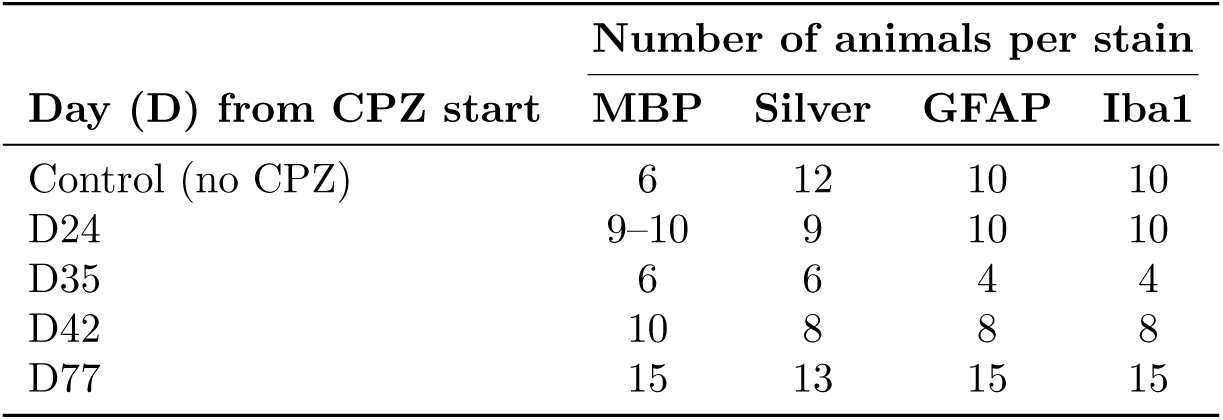
Breakdown of animals (all males) used for histological evaluation for each stain. MBP - Myelin Basic Protein; GFAP – Glial Fibrillary Acidic Protein; Iba1 – Ionized calcium-binding adaptor molecule 1.

### Number of animals per MRI study

The animals were group housed (2-5 per cage) and maintained on a 12:12 hour light–dark cycle (lights on at 7am). Food and water were available ad libitum. Animals were kept in the local facility for minimum of 7 days prior to experimentation and were weighed at least twice weekly. Animals were brought to the MRI suite a minimum of 1 day prior to in vivo MR imaging and spectroscopy (MRI/S) acquisition to acclimatise.

### Number of animals per stain

#### Study design

A schematic of the study design is shown in Figure1. For the in vivo study, the animals were pooled from three in vivo experiments, where one group of mice received a vehicle treatment, and the others did not receive any vehicle treatment. The imaging protocol was identical for all animals and time-points. All animals were imaged at baseline (pre-cuprizone), and several times post CPZ (Days 24, 35, 49, 63, 77 from CPZ start; see Figure 1).

**Figure 1:**
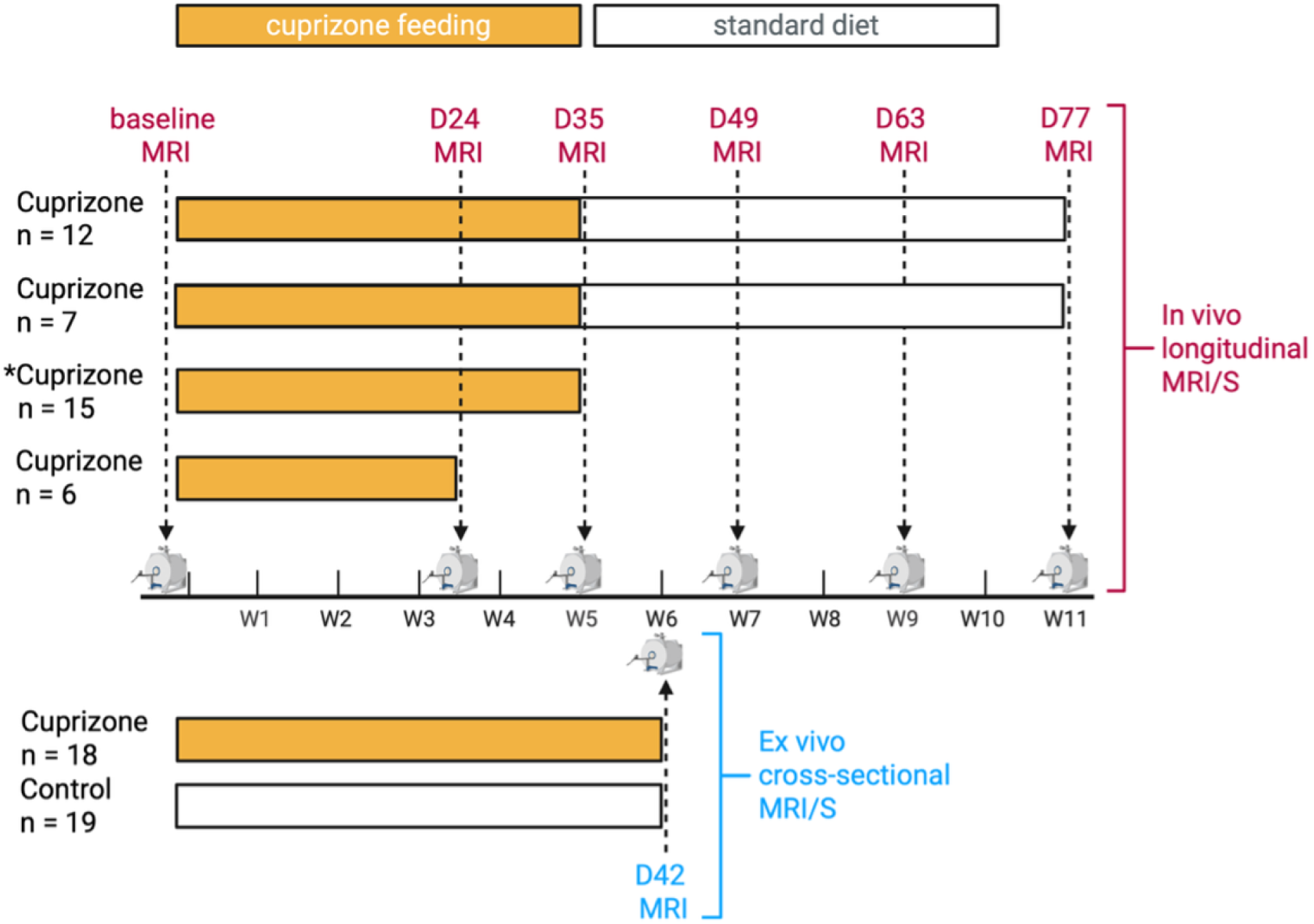
Study design. Schematic representation of the in vivo and ex vivo study designs. Scans are shown by the broken line in foreground.

As there were no differences in brain volumes due to time or cohort (see Figure S5) we pooled the data from all mice and conducted an analysis that compared each post-cuprizone timepoint to a pre-cuprizone baseline with an additional within-subject modelling. These three in vivo studies took place from May 2022 to April 2023. To enable histological evaluation, a group of animals randomly pre-selected before the start of the study were sacrificed on days 24, 35, 42, and 77 (see Table 2). The ex vivo study was only one, taking place between Jan 2022 to April 2022 (see Table 1).

#### In vivo study

Cuprizone (Bis(cyclohexanone)oxaldihydrazone; C9012, Sigma Aldrich) was administered to all mice (n = 40) for 35 consecutive days as 0.2% cuprizone mixed into standard powdered rodent food provided ad libitum, as described previously (Wood et al., 2016). Cuprizone treatment was stopped at day 35. Some mice (n = 15) within the in vivo study separately served as the control arm of a drug study. These mice received vehicle (0.5% hydroxypropyl methyl cellulose, 0.02% Tween-80 and 99.48% H2O) daily at 10ml/kg by oral gavage in conjunction with cuprizone treatment.

For all mice, baseline MRI/S were acquired 1-3 days before commencement of cuprizone administration. MRI/S were longitudinally acquired on day 24 only (n = 6), on days 24 and 35 (n = 15), on days 35, 49 and 77 (n = 7), and on days 24, 35, 49, and 77 (n = 12). After the final MRI/S acquisition session, mice were immediately sacrificed, and brain tissue was collected for histology.

#### Ex vivo study

Cuprizone (Bis(cyclohexanone)oxaldihydrazone; C9012, Sigma Aldrich) was administered to all mice (n = 37) for 42 consecutive days as 0.2% cuprizone mixed into standard powdered rodent food provided ad libitum, as described previously (Wood et al., 2016). After 42 days, mice were sacrificed and their heads processed for ex vivo MRI acquisition. A group of 16 mice within the ex vivo study additionally served as the control arm of a genetics study (reported elsewhere, manuscript submitted) and as such received 125mg/kg tamoxifen by intraperitoneal injection daily for 4 consecutive days – this treatment did not affect any baseline changes in their brain images (data not shown). Tamoxifen treatment was stopped at least 21 days before beginning cuprizone treatment.

### MRI/S acquisition

#### In vivo study

Mice were anaesthetised with 5% isoflurane in oxygen:air (30:70) for approximately 3 minutes, after which isoflurane was lowered to 2% and the mouse placed inside scanner bed and radiofrequency coil. MRI/S were acquired using a 9.4T Bruker small animal scanner with an 86 mm (diameter) transmit volume coil and a mouse brain 2×2 surface array receiver coil.

Diffusion tensor imaging (DTI) data were acquired with a single-shot SE-EPI (spin-echo echo planar imaging) using the following parameters: *T_E_* = 20.4 ms, *T_R_* = 3000 ms, FOV = 18×12.15 mm, matrix = 80×54 (partial-FT 1.6 in the phase-encoding direction), slice thickness = 0.4 mm, 30 contiguous slices, five b0 images, 30 diffusion weighting directions, b-value = 1000 s/mm^2^, *δ*_Δ_ = 3/11 ms, bandwidth = 357 kHz. Transmit *B*_1_ Actual Flip Angle Imaging (TB1AFI) data were acquired using the following parameters: *T_E_*= 2.85 ms, *T_R_*_1_ = 20 ms, *T_R_*_2_ = 100 ms, 1 average, flip angle = 55°, matrix 40×40×24, FOV = 16×16×9 mm, slice thickness = 9 mm, bandwidth = 25 kHz.

Each Multiparametric mapping (MPM) scan with 150×150×150 µm resolution was comprised of three different 3D acquisitions designed to provide magnetization-transfer-weighted (MTw), proton-density-weighted (PDw), and T1-weighted (*T*_1_w) images using the following parameters: MT pulse shape = gaussian, MT pulse length = 4 ms, MT pulse bandwidth = 685 Hz, MT pulse offset = -3000 Hz, FOV = 16×16×9 mm, matrix = 108×108×60 (partial-FT 1.33 in both phase-encoding directions), 2 averages, bandwidth = 100 kHz. Other specific parameters are reported in Table 3 and representative MPM images are shown in Figure S2.

**Table 3:** Multiparametric mapping (MPM) acquisition parameters.

| | PDw | $T_1w$ | MTw |
| --- | --- | --- | --- |
| $T_R$ (ms) | 20 | 20 | 24 |
| Flip angle ($^\circ$) | 4 | 20 | 6 |
| $T_{E1}$ (ms) | 2.5 | 2.5 | 2.5 |
| $T_E$ spacing (ms) | 2.1 | 2.1 | 2.1 |
| Number of echoes | 7 | 7 | 6 |

Single voxel ^1^H-MRS was acquired over the corpus callosum (CC) area (Figure S4). A Point REsolved Spectroscopy (PRESS) pulse sequence was used with the following parameters: *T_E_*= 8.4 ms, *T_R_*= 2500 ms, 720 averages, acquisition bandwidth = 4400 Hz, 2048 acquisition points, voxel size = 2.5×0.75×3 mm. Outer volume suppression and water suppression with variable pulse power and optimized relaxation delays (VAPOR) were used to mitigate the contribution of signal from outside the prescribed voxel and suppress unwanted signal from water.

#### Ex vivo study

Mice from the ex vivo study were killed by transcardiac perfusion with room temperature (RT) heparinized PBS followed by 10% formalin. After perfusion-fixation, mice were decapitated and their heads maintained in 10% formalin for 24 hours at RT on a rotator. Heads were then rehydrated in PBS with 0.05% sodium azide for minimum 28 days at 4°C.

MRI was acquired using a 9.4T Bruker small animal scanner. Heads were immersed in fluorinated liquid (Galden, Solvay) to reduce susceptibility artefacts and loaded into a 39mm quadrature volume coil. T2-weighted RARE images were acquired using the following parameters: RARE factor = 8, *T_E_* = 30 ms, *T_R_* = 3000 ms, FOV = 25×25×20 mm, matrix = 250×250×200, bandwidth = 55 kHz, scan time = 5 hours 44 minutes.

Magnetization-Prepared 2 Rapid Gradient-Echo (MP2RAGE) (Marques et al., 2010) was used to acquire images at two inversion times (*T_I_*): *T_I_*_1_ = 800 ms, *T_I_*_2_ = 3200 ms, flip angle = 7°, *T_E_*= 3.2 ms, *T_R_* = 8.36 ms, segment *T_R_* = 6000 ms, FOV = 25×25×20 mm, matrix = 200×200×160, bandwidth = 49 kHz, 15 averages, scan time = 4 hours 24 minutes.

DTI data were acquired with a 2D multi-slice, standard spin echo acquisition: *T_E_*= 22.6 ms, *T_R_*= 4000 ms, FOV = 25.6×25.6, matrix = 128×128, 67 slices with 0.2-mm thickness and 0.1-mm gap, 4 b0 images, 30 diffusion weighting directions, b-value = 1500 s/mm^2^, *δ*_Δ_ = 4/10 ms, bandwidth = 20 kHz, scan time = 3 hours 38 minutes.

### MRI and MRS analyses

All images were automatically converted to NIfTI format using BrkRaw (Lee, 2020).

#### MPM processing

The processing workflow for MPM images was optimised for this study, as detailed below. TB1AFI, *T*_1_w and MTw images were registered to PDw images with rigid body transformations using Advanced Normalization Tools (ANTs) (Klein et al., 2009). MPM images from the baseline scan were co-registered across subjects and averaged to create a study template using the ANTs script antsMultivariateTemplateConstruction2.sh. All images acquired in this study were non-linearly registered to this template using antsRegistration from ANTs in a separate stage. The Jacobian determinant map of each subject’s transformation to this space was subsequently used to calculate the voxelwise volume difference maps, both accounting for (absolute) and discounting (relative) individual variation in total brain volume. Additionally, the study template was normalised to the Allen Mouse Brain Common Coordinate Framework (CCFv3), facilitating the use of the Allen Brain Atlas (see ROI analysis section).

R2* and PDw-, *T*_1_w-, and MTw-S0 maps were calculated with the ECSTATICS method (Weiskopf et al., 2014) using the qi mpm_r2s command in QUantitative Imaging Tools (QUIT) (Hawkins et al., 2018). B1 maps were calculated from TB1AFI images using the qi afi command in QUIT. The S0 and B1 maps were used to calculate maps of MTsat*δ* and longitudinal relaxation rates (*R*_1_) with the qi mtsat command in QUIT. Representative MPM maps are shown in Figure S2. Figure S3 additionally shows an overlay of MTsat*δ* with a myelin histological stain, demonstrating close correspondence between the two measures.

#### DTI

DTI images were first denoised and deringed using MRtrix3 toolbox (Tournier et al., 2019), distortion corrected using FSL’s topup (Andersson, 2003) and corrected for eddy currents and movements using FSL’s eddy (Andersson, 2016). The denoised and motion/distortion corrected images were then registered to the MTw image using antsRegistration from ANTs (Klein et al., 2009). The inverse transformation derived from a given subject’s registration to the study template was applied to the template brain mask to mask the DWI images. Finally, fractional anisotropy (FA) and diffusivity (mean, radial and axial) maps were calculated using the dtifit function in FSL’s FDT toolbox. These maps were then transformed to the study template space using the previously calculated forward transformation.

#### MRS

The following molecules were measured using the PRESS ^1^H-MRS technique: *γ*-aminobutyric acid (GABA), glutamine (Gln), glutamate (Glu), glutathione (GSH), inositol (Ins), N-acetylaspartate (NAA), taurine (Tau), glycerophosphochoine + phosphocholine (GPC+PCh), creatine + phoshphocreatine (Cr+PCr), glutamate + glutamine (Glu+Gln). MRS raw data were analysed using a custom Matlab® pipeline. MR spectra were then analysed with two software packages: FID Appliance (FID-A) and Linear Combination (LC) Model version 6.3. First, FID-A was used to pre-process ^1^H-MRS data, simulate the metabolites, and create a basis set (model spectra). Then, we used LCModel to calculate the water-referenced concentration (in mM) of the different metabolites by applying linear combinations of the model spectra to determine the best fit of the individual ^1^H-MRS data. Finally, the method of Cramér Rao (Cramér Rao Lower Bound, CRLB) was applied to ensure the reliability of the metabolite quantification, by which metabolite concentrations with S.D. *≥* 20% are classified as not accurately detectable and are discarded. MRS voxel position and a representative MR spectrum are shown in Figure S4.

#### MP2RAGE

The MP2RAGE images were processed using QUIT (Hawkins et al., 2018). The complex image from each coil channel was combined using the UTE reference image and the COMPOSER method (Robinson et al., 2017). The combined image was input to qi_mp2rage, which uses the signal at both inversion times to produce a *T*_1_ map and a *T*_1_w image that is inherently corrected for B1 field inhomogeneity, thus addressing non-uniformity of the signal intensity across an image.

### Statistical analysis

#### Voxelwise analysis

MPM and DTI maps were analysed using voxel-based statistical parametric map comparison and Region-Of-Interest (ROI) analysis. For each time point, within-subject longitudinal differences from baseline were calculated for Jacobian determinant, MTsat*δ*, FA, and MD maps. One-sample (change from baseline) and two-sample (comparison between drug groups) tests were performed on these difference maps using voxelwise permutation tests using FSL randomise with 5000 permutations, threshold-free cluster enhancement, and controlling for family-wise error (FWE) rate (Winkler et al., 2014). The results are visualized using the dual-coding approach (Allen et al., 2012; Taylor et al., 2023): the effect size, i.e. the magnitude of the differences, is represented by the colour of the overlay. The overlay transparency increases with the p value derived from the statistical parametric maps. Additionally, black contour lines denote clusters that surpass a significance threshold of FWE-corrected p *<* 0.05.

#### MRS analysis

Changes in MRS metabolites from baseline were analysed using R (v4.4.2). First, the metabolites were z-scored using each cohort’s baseline mean and standard deviation for each metabolite. Three subjects (2 from DD36 at day 49, and 1 from DD36 at day 77) were excluded on the basis that all their normalised metabolite values were Z *>* 10, as this was believed to indicate technical issues with the spectra. A mixed-effects model was fit to predict z score with a random intercept per subject, while metabolite and the interaction between metabolite and session were provided as fixed effect predictors, using lme4 (Bates, 2015) (version 1.1-35.5) and lmerTest (Kuznetsova, 2017) (version 3.1-3). The estimated marginal means (EMM) were calculated from model fit with emmeans (v 1.10.5), and 95% confidence intervals are shown in Figure 8, adjusted with Šidák’s correction to account for the 10 metabolite comparisons within a session. An EMM interval that did not include 0 was considered to be significantly different from baseline.

#### ROI analysis

Relevant metrics were extracted from volume, diffusion, and relaxometry maps from 16 regions, derived from the Allen Mouse Brain Common Coordinate Framework (Wang et al., 2020), spanning grey and white matter in the cerebrum and cerebellum. Regional values for each MRI metric were normalised using the mean and standard deviation of the respective values for each region from the baseline scans. A mixed-effects model was fit for each metric, with a random intercept per subject, and session and region as fixed effect predictors, with lme4 (Bates, 2015) (version 1.1-35.5) and lmerTest (Kuznetsova, 2017) (version 3.1-3) in R (v4.4.2). The EMMs were calculated from each model fit with emmeans (v 1.10.5), and 95% confidence intervals are shown in Figure 6. Šidák’s correction was applied to account for the 16 regional comparisons within a session. An EMM interval that did not include 0 was considered to be significantly different from baseline.

### Histological markers and tissue processing

Mice from the in vivo study (Table 2) were killed by transcardiac perfusion with heparinised sterile saline at RT. After perfusion, whole brains were dissected. The left hemisphere was immersion-fixed in 4% paraformaldehyde for 48-72 hours at 4°C, and cryoprotected thereafter in 30% sucrose at 4°C for at least an additional 72 hours. After cryoprotection, left hemisphere samples were stored at 4°C until sectioned using a freezing microtome. For each left hemisphere, 20µm thick sections were cut in 12 series.

Tissue staining involved histochemistry with Bielschowsky Silver staining to visualize nerve fibres, immunohistochemistry (IHC) with markers of astrocytes (GFAP; glial fibrillary acidic protein), microglia (Iba1; Ionized calcium binding adaptor molecule 1) and immunofluorescence for myelin basic protein (MBP) to further characterise axonal pathology.

#### Bielschowsky silver staining

Bielschowsky silver staining was used to visualise nerve fibres in the tissue, enabling the quantification of myelination (Pfeifenbring et al., 2015; Preziosa et al., 2019). Free-floating left hemisphere sections were washed in Tris-buffered saline (TBS) and slide-mounted to dry overnight. Slides were then washed three times in distilled water and incubated in 10% silver nitrate solution at 40°C for 15 minutes until sections became a light brown colour. Slides were then removed and concentrated ammonium hydroxide was added dropwise until clearing. Cleared slides were incubated in ammonium silver solution at 40°C for 30 minutes. Slides were then placed in developer working solution (prepared as follows: 8 drops of concentrated ammonium hydroxide and 8 drops of developer stock solution [100ml distilled water, 20ml formaldehyde 37-40%, 0.5g citric acid and 2 drops nitric acid in 50ml of distilled water] for 1 minute, 1% ammonium hydroxide solution for 1 minute, 5% sodium thiosulfate solution for 5 minutes, and then washed three times with distilled water. Slides were then dehydrated and cleared through 70%, 90%, 2x 100% IMS and 2x xylene, and a coverslip applied with DPX (06522, Sigma-Aldrich).

#### 3, 3’-diaminobenzidine (DAB) staining

Free-floating left hemisphere sections were washed in TBS before being heated to 80°C in 10 mM sodium citrate buffer (pH 6.0) for 30 minutes for antigen retrieval. Sections were then washed again in TBS before incubation in 1% hydrogen peroxide (H_2_O_2_) for 15 minutes. Sections were washed in TBS before incubation in TBS with 0.2% Triton-X and 10% skimmed milk powder (SMP) for 40 minutes to block non-specific binding. Sections were incubated overnight at 4°C with primary antibodies against either Iba1 (rabbit anti-Iba1; 1:2500, 019-19741, Alpha Laboratories) or GFAP (rabbit anti-GFAP; 1:2000, Z033401-2, Agilent Technologies). Sections were then washed in TBS and incubated in secondary antibody (biotinylated goat anti-rabbit; 1:1000, BA-1000, Vector Laboratories Ltd.) for 2 hours, before being washed and incubated in Vectastain ABC Kit (PK-6000, Vector Laboratories Ltd.) for 1 hour on a shaker at RT. Staining was performed by incubation in DAB solution (1 DAB tablet [J60972.KW, Alfa Aesar] per 10ml TBS with 3µl H_2_O_2_). Finally, sections were mounted onto slides and left overnight. They were then dehydrated and cleared through 70%, 90%, 2x100% IMS and 2x xylene, and a coverslip applied with DPX.

#### Immunofluorescence staining

Free-floating left hemisphere sections were washed in TBS before being heated to 80°C in 10mM sodium citrate buffer (pH 6.0) for 30 minutes for antigen retrieval. Sections were then washed again in TBS, before incubation in TBS with 10% normal horse serum and 0.2% Triton-X for 40 minutes to block non-specific binding. After this, sections were incubated overnight at 4°C with primary antibody against MBP (rabbit anti-MBP, 1:5000, AB7349, Abcam). Sections were then washed in TBS and incubated in secondary antibody (Alexa-Fluor 647 donkey anti-rabbit, 1:1000, A31573, ThermoFisher Scientific) for 2 hours at RT. Finally, sections were counterstained with 1µg/mL DAPI and coverslipped with FluorSave (345789, Sigma-Aldrich).

### Histological image acquisition and analysis

Iba1, GFAP and silver-stained sections were imaged using Olympus VS120 Slide Scanner (Olympus, Japan) at 40X magnification. MBP stained sections were imaged using Olympus VS300 Slide Scanner (Olympus, Japan) at 20X magnification. The scanned images were processed in Olympus OlyVIA (v2.9). Snapshots of stained sections containing the ROIs were taken at 4X magnification (Iba1, GFAP, MBP) or 1X magnification (silver).

OlyVIA snapshots were then exported to ImageJ for analysis. Iba1 and GFAP images were converted to 8-bit colour, ROIs were selected, and the thresholds were adjusted. For all stains, at least 4 images for each ROI of each mouse were analysed and averaged to avoid bias selection.

Iba1 and GFAP staining were quantified in ImageJ by thresholding and calculating the area fraction of positive signal within each ROI. MBP and silver staining were quantified as mean grey values, normalized to background. Statistical analysis was performed with GraphPad Prism (v 10.6.0) using mixed ANOVA, with region as the within-subject factor and timepoint as the between-subject factor. Post-hoc comparisons of each cuprizone group versus control were conducted using Dunnett’s test.

## Results

### Myelination changes persist 5 weeks after cuprizone cessation

#### MTsat*δ*

In vivo MTsat*δ* images, as expected (Henkelman et al., 2001; Helms et al., 2008), provided clear grey-white matter contrast, confirming MTsat*δ* as a sensitive parameter for detecting regional myelination changes. Voxelwise clusters of mean regional differences in MTsat*δ* (Figure 2) showed significant reductions by Day 24 in the corpus callosum and deep cerebellar nuclei, suggesting substantial myelin loss. By Day 35, demyelination expanded into cortical regions, striatum, and nucleus accumbens. Only partial recovery was evident by day 77 (5 weeks after cessation of cuprizone treatment) and many abnormalities remained. Atlas-based regional analysis (Figure 6) revealed early decreases by Day 24 in white matter regions of the corpus callosum, fornix and cerebellar arbor vitae; the latter was significantly reduced only at Day 24, whereas the others remained reduced at least through Day 63. Interestingly, many grey matter regions exhibited widespread MTsat*δ* reductions, with the largest changes in the cerebellar nuclei, and additional decreases in the isocortex and striatum. Smaller, transient decreases were also observed in the olfactory areas, pallidum and septum. At the final time point (Day 77) significant reductions remained in the cerebellar nuclei, isocortex, striatum and corpus callosum.

**Figure 2:**
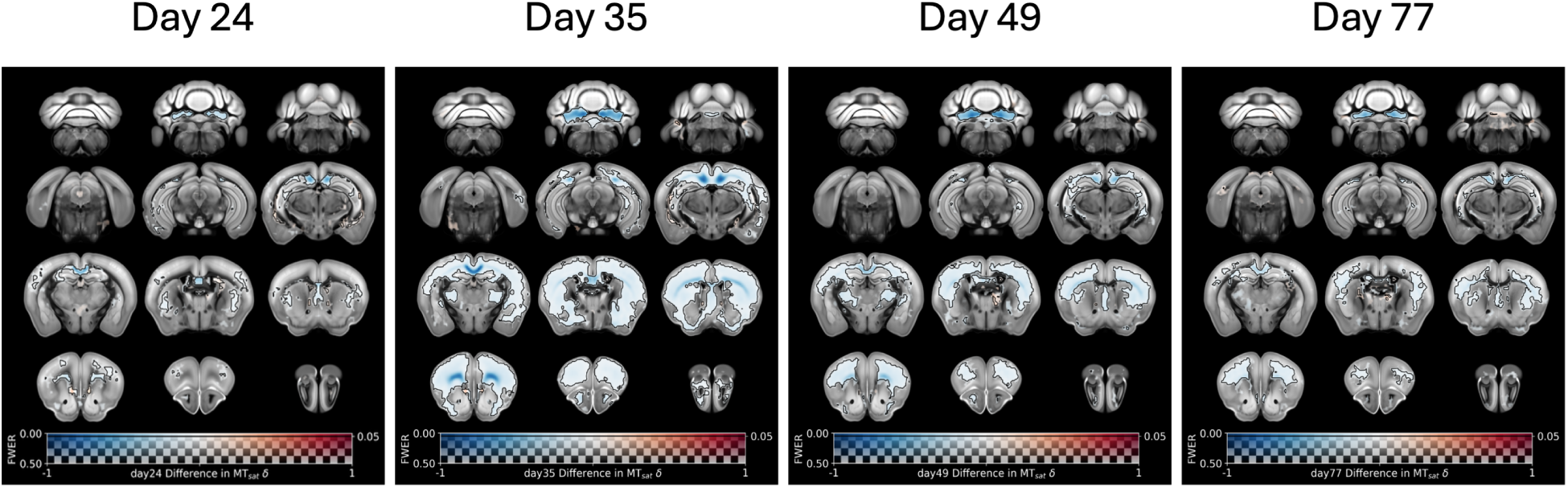
Time evolution of mean differences in MTsat*δ*. Longitudinal voxelwise comparison of MTsat*δ* at four timepoints versus pre-cuprizone baseline: difference is shown overlaid on a template, where the magnitude of the difference is coded into colour. Transparency is controlled by statistical significance (p value, where small p-values are more opaque and values above 0.5 are completely transparent). P-values were generated using TFCE on the contrast for a two-tailed, two sample t-test. Black contours highlight voxel clusters where FWE-corrected p *<* 0.05.

#### Relaxation rate (*R*_1_)

Areas of voxelwise R1 differences (Figure 3) partially corroborated the MTsat*δ* findings, although *R*_1_ changes were smaller in magnitude and more focal than the widespread MTsat*δ* effects. Reductions in *R*_1_ were detected primarily in the corpus callosum at Day 24; by Day 35 they were also evident in the deep cerebellar nuclei, and by Day 49 they had extended to the cortex and hippocampus. These changes showed only partial recovery by day 77. Regional analysis (Figure 6) of white matter areas revealed sustained *R*_1_ reductions in the corpus callosum ROI from Day 24 through Day 77, peaking at Day 35, while the fornix exhibited partial reductions during the later phase (Days 49 – 63). In grey matter, *R*_1_ decreased in the cerebellum - transient and early in the cortex, robust and persistent in the deep nuclei. Additional late phase, small reductions were observed in the pallidum and thalamus (Days 63-77).

**Figure 3:**
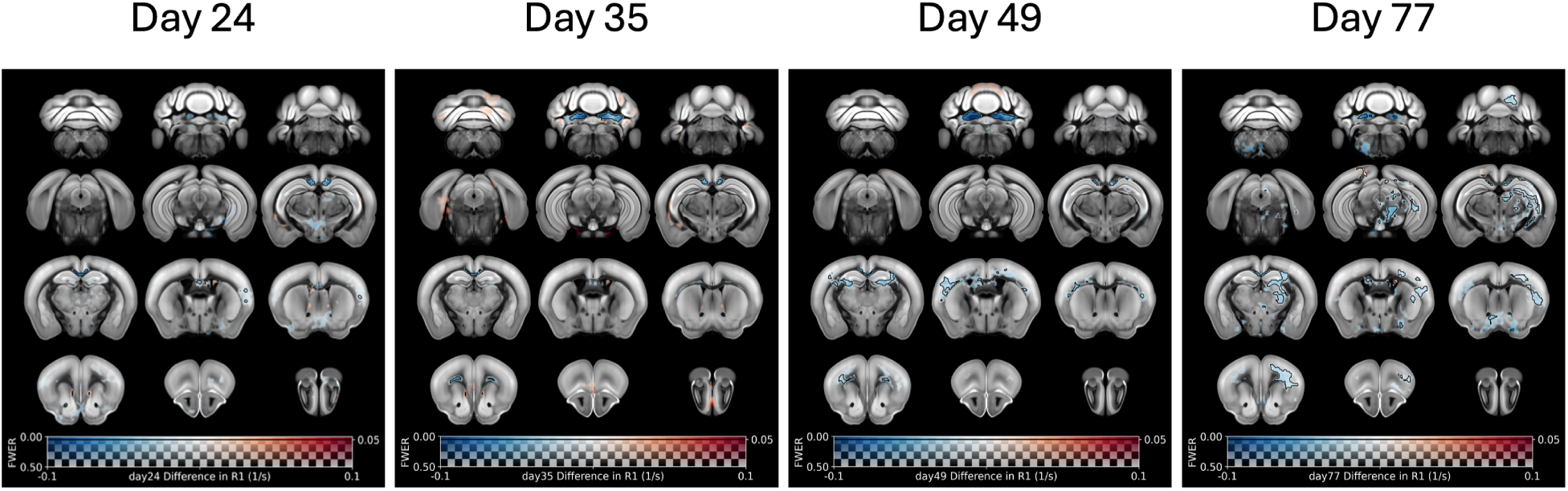
Time evolution of mean differences in *R*_1_. Longitudinal voxelwise comparison of *R*_1_ at four timepoints versus pre-cuprizone baseline: difference is shown overlaid on a template, where the magnitude of the difference is coded into colour. Transparency is controlled by statistical significance (p value, where small p-values are more opaque and values above 0.5 are completely transparent). P-values were generated using TFCE on the contrast for a two-tailed, two sample t-test. Black contours highlight voxel clusters where FWE-corrected p *<* 0.05.

### Longitudinal DTI can track cuprizone-induced microstructural disruption and spontaneous repair

Mice also underwent diffusion tensor imaging (DTI) to derive fractional anisotropy (FA), mean diffusivity (MD), axial diffusivity (AD), and radial diffusivity (RD). Voxelwise and atlas-based regional changes from baseline (Day 0) are shown in Figure 4 and Figure 6, respectively.

**Figure 4:**
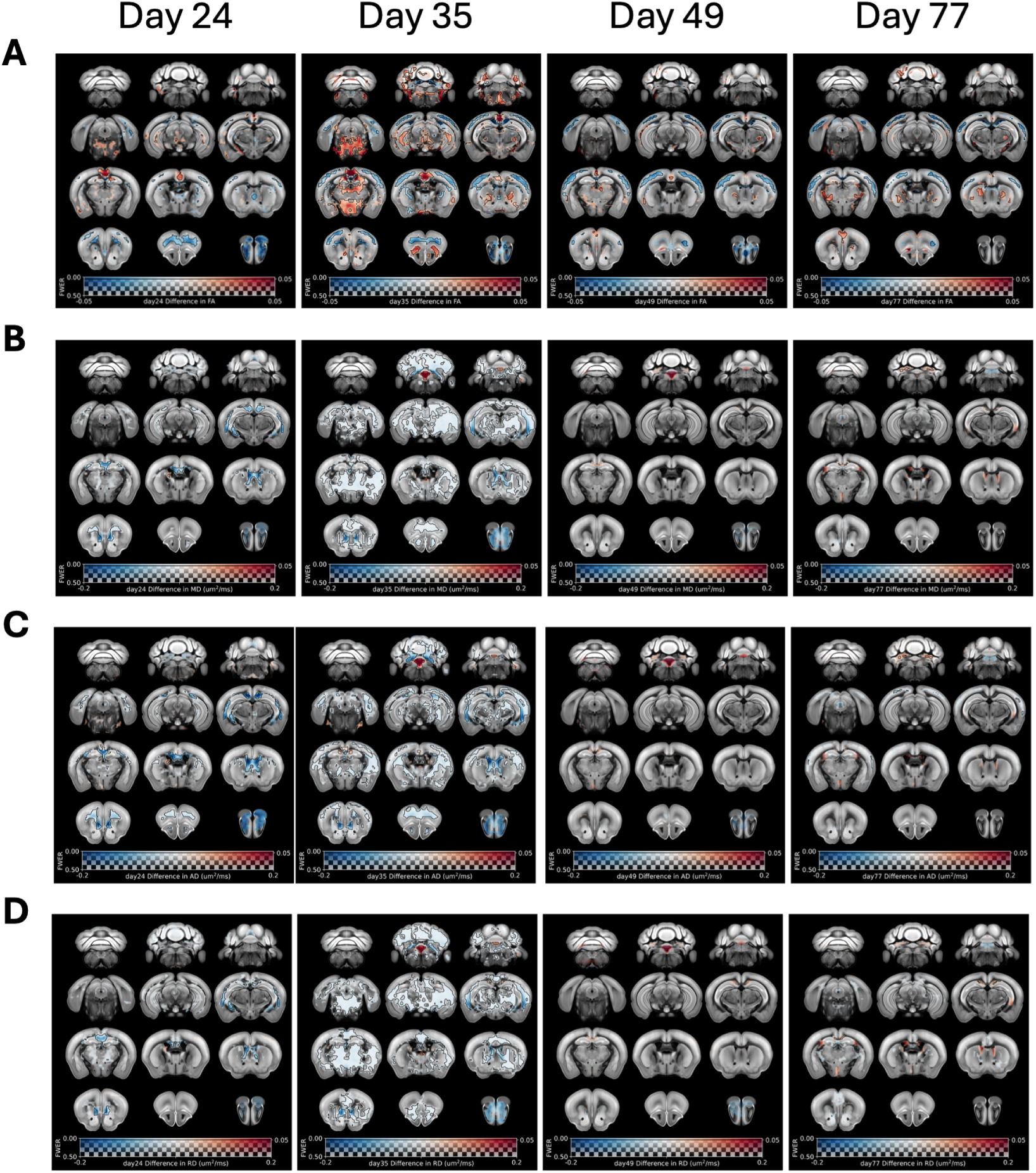
Time evolution of mean changes in A. fractional anisotropy (FA), B. mean diffusivity (MD), C. axial diffusivity (AD), and D. radial diffusivity (RD). Longitudinal voxelwise comparison of DTI metrics at four timepoints versus pre-cuprizone baseline: difference is shown overlaid on a template, where the magnitude of the difference is coded into colour. Transparency is controlled by statistical significance (p value, where small p-values are more opaque and values above 0.5 are completely transparent). P-values were generated using TFCE on the contrast for a two-tailed, two sample t-test. Black contours highlight voxel clusters where FWE-corrected p *<* 0.05.

FA showed a mix of increases and decreases with a focal pattern apparently aligned to anatomical boundaries (Figure 4A and Figure 6). In white matter, FA was predominantly increased in the cerebellum and internal capsule, with no significant elevation at the earliest timepoint (Day 24) but rising thereafter and remaining elevated through Day 77. A slight, transient FA increase was also observed in the optic and other fibre tracts between Days 35–49. In grey matter, FA decreased across large portions of the cerebral cortex (“isocortex” ROI) up to and including Day 77, while small, well-delineated FA increases appeared in parts of the cingulate and retrosplenial cortices during the early phase (Days 24–35). Additional FA increases were evident in the brainstem mostly at Days 24–35. FA was also increased in subcortical grey matter of the thalamus (D35–D77), with an emerging increase in the striatum at D77. Qualitatively, the largest FA changes were observed at D35, and many persisted without full resolution by Day 77.

MD, by contrast, was less affected, showing smaller clusters of change, many of which resolved by Day 77 (Figure 4B and Figure 6). In white matter, MD exhibited early decreases across several regions, including the corpus callosum and internal capsule; aside from a small increase in the internal capsule, these were not significant at Day 77. In grey matter, MD was similarly decreased during the early phase in the cerebellum, cortex, and subcortex. Notably, MD in the cerebellar nuclei switched from an initial decrease to an increase between Days 35 and 49; a similar pattern (early decrease with a later upward trend) was seen in select white-matter regions (fornix and optic nerve), although these latter changes did not reach significance.

AD and RD were not examined in detail for regional trajectories. Their voxelwise differences from baseline are shown in Figure 4C–D. Overall, both metrics exhibit a mixed pattern of changes that broadly parallels the MD findings, as expected.

### Cuprizone treatment results in prolonged hippocampal, cortical and cerebellar volumetric changes

Regional brain volumes, assessed by tensor-based morphometry (TBM) also showed widespread time-related changes (Figure 5 and Figure 6). At Day 24, there were significant increases in the volume of the hippocampus (shown only in Figure 5) and deep cerebellar nuclei, with significant decreases also present, particularly in cortical and subcortical regions, and the cerebellar cortex. By Day 35, areas of change were more clearly defined, and volume changes more pronounced. Although some reversal was evident by Day 49, changes in the hippocampus, cerebellum and cortex persisted to Day 77.

**Figure 5:**
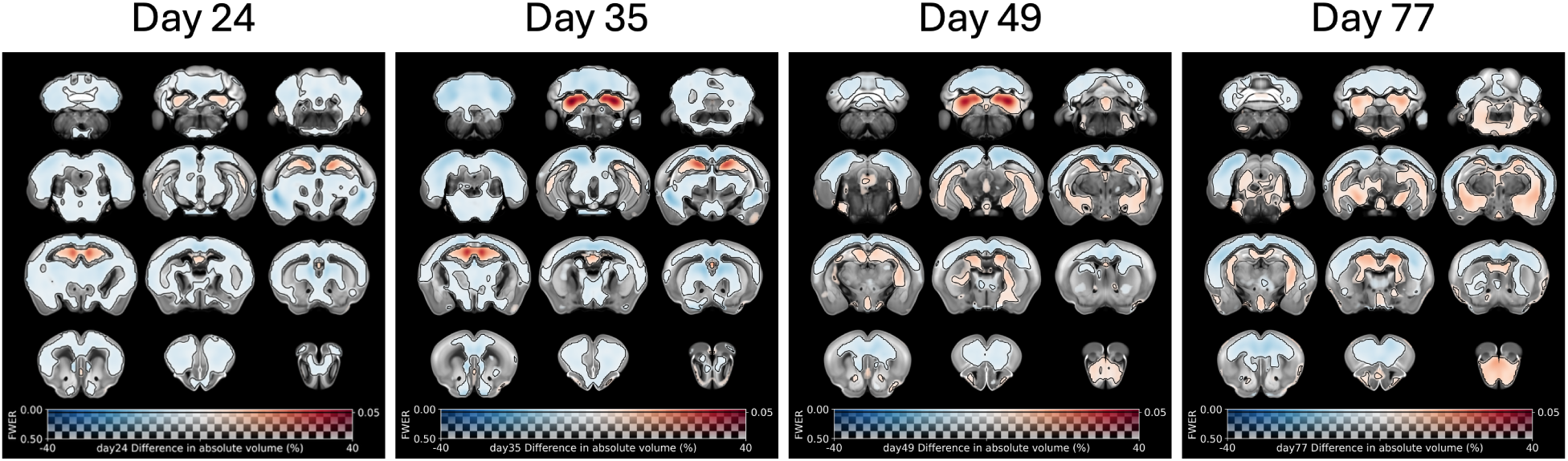
Time evolution of regional volume changes. Longitudinal voxelwise comparison of absolute volume at four timepoints versus pre-cuprizone baseline: difference is shown overlaid on a template, where the magnitude of the difference is coded into colour. Transparency is controlled by statistical significance (p value, where small p-values are more opaque and values above 0.5 are completely transparent). P-values were generated using TFCE on the contrast for a two-tailed, two sample t-test. Black contours highlight voxel clusters where FWE-corrected p *<* 0.05.

### Regions of interest (ROIs)

As all images were registered to common space, we performed atlas-based analysis by extracting the mean values from all metrics used in the structural imaging in this study. The 16 ROIs were derived from the Allen brain atlas, and the results have been discussed in the relevant sections above.

### Ex vivo structural imaging partially corroborates in vivo results

A cohort of mice that did not undergo in vivo imaging was sacrificed on Day 42, immediately after cessation of cuprizone treatment; their brains were imaged ex vivo and compared with those of control mice maintained under identical conditions but not exposed to cuprizone (see Figure 1 for the experimental design). The ex vivo metrics analysed were the *R*_1_ relaxation rate (1*/T*_1_), FA, MD, and TBM-derived regional volumes, all presented as voxelwise differences in Figure 7.

**Figure 6:**
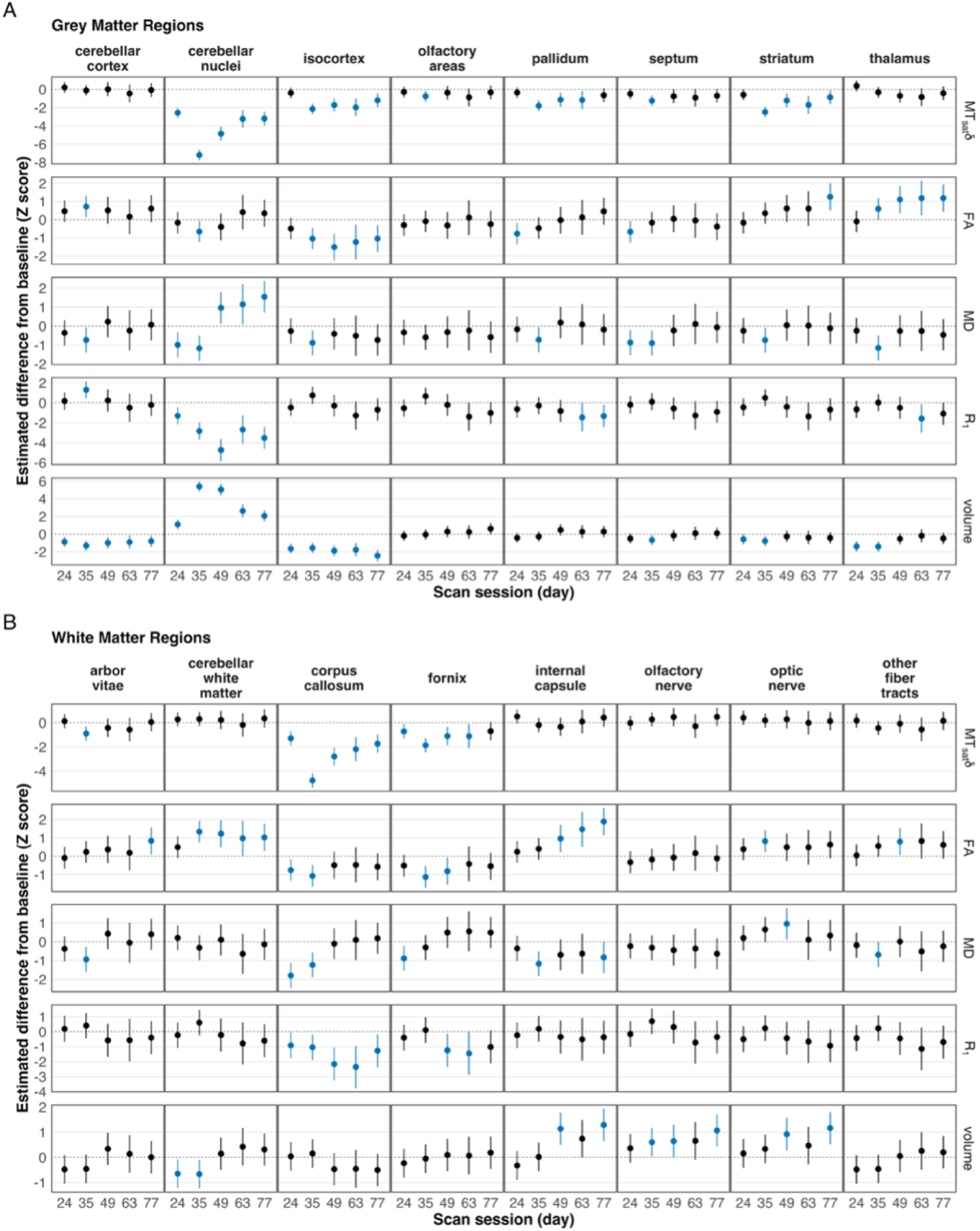
Regional changes from baseline in MRI metrics (MTsat*δ*, fractional anisotropy (FA), mean diffusivity (MD), R1 and volume) in A. grey matter ROIs and B. white matter ROIs. Regional values for each MRI metric were normalised using the mean and standard deviation of the respective values for each region from the baseline scans. A mixed-effects model was fit for each metric, with a random intercept per subject, and session and region as fixed effect predictors. The estimated marginal means (EMM) were calculated from each model fit, and 95% confidence intervals are shown. Šidák’s correction was applied to account for the 16 regional comparisons within a session. An EMM interval that did not include 0 was considered to be significantly different from baseline (shown in blue).

**Figure 7:**
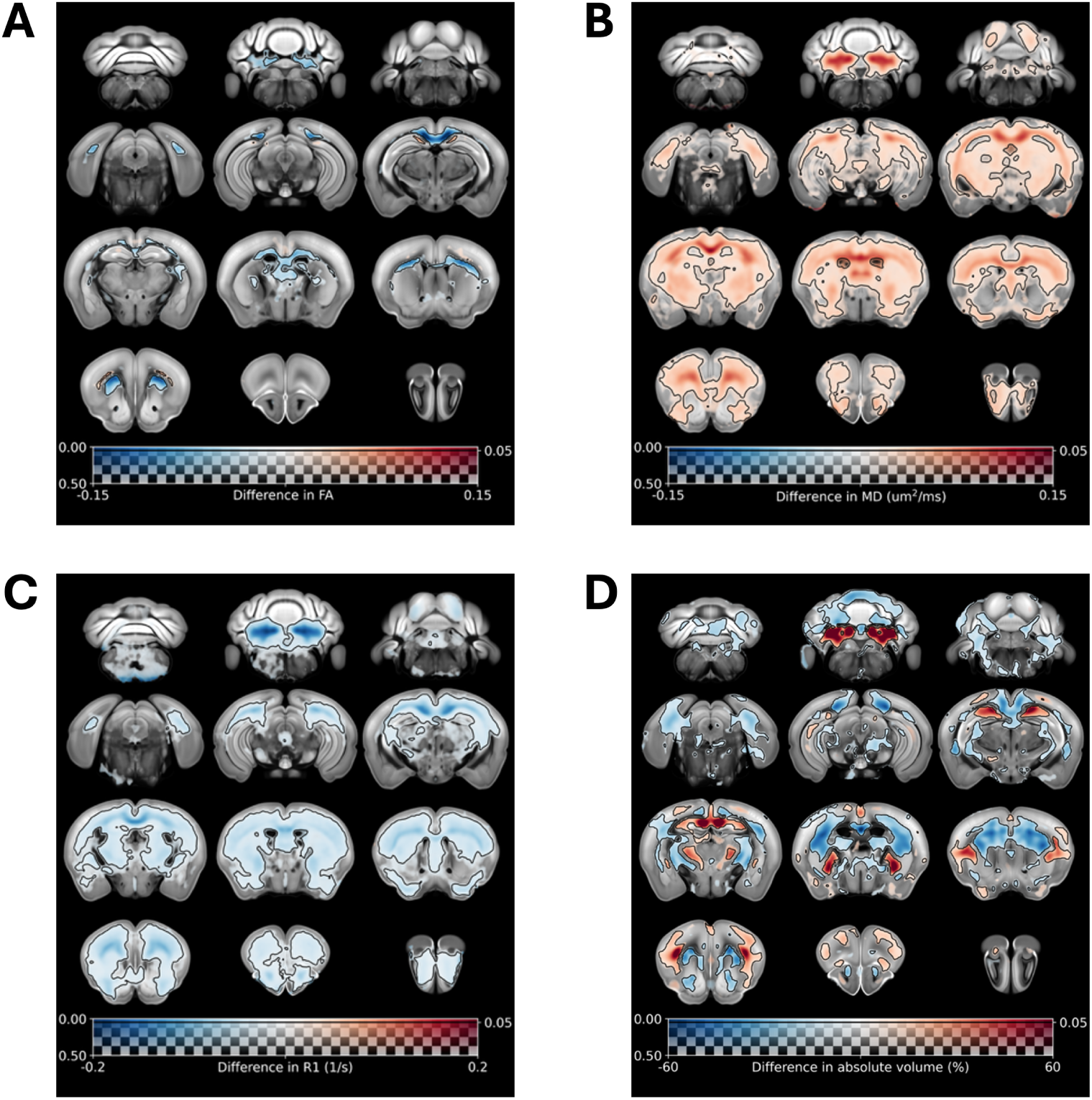
Ex vivo changes in A. fractional anisotropy (FA), B. mean diffusivity (MD), C. *R*_1_ (1*/T*_1_), and D. volumes (tensor based morphometry, TBM) in cuprizone-treated compared to control mice at day 42. Cross-sectional voxelwise comparison of DTI, *R*_1_ and TBM metrics immediately after cuprizone cessation at Day 42 versus no cuprizone control mice: differences are shown overlaid on a template, where the magnitude of the difference is coded into colour. Transparency is controlled by statistical significance (p value, where small p-values are more opaque and values above 0.5 are completely transparent). P-values were generated using TFCE on the contrast for a two-tailed, two sample t-test. Black contours highlight voxel clusters where FWE-corrected p *<* 0.05.

At Day 42, ex vivo FA (Figure 7A) showed focal decreases predominantly within the corpus callosum. Ex vivo MD (Figure 7B) was broadly elevated across white matter, particularly in the corpus callosum, and extended into cortical and subcortical regions including thalamus, cortex, hippocampus, and the deep cerebellar nuclei, with smaller effects in the cerebellar cortex. *R*_1_ (Figure 7C) was decreased in multiple areas, notably the deep cerebellar nuclei, corpus callosum, hippocampus, and cortex. The pattern of *R*_1_ decreases (qualitatively) resembled the changes in MTsat*δ* seen in in vivo images at both Day 35 and 49 (Figure 2).

TBM (Figure 7D) revealed regional volume changes consistent with the in vivo findings at Days 35 and 49: increases in the deep cerebellar nuclei and hippocampus, alongside decreases in the cerebellum and cortico-striatal regions. Cortical areas showed a mixture of increases and decreases, but did not recapitulate the prominent cortical decreases observed in vivo.

### MRS captures widespread metabolic alterations in the corpus callosum

We also performed ^1^H-MRS in the corpus callosum (Figure S4), quantifying *γ*-aminobutyric acid (GABA), glutamine (Gln), glutamate (Glu), glutathione (GSH), myo-inositol (Ins), N-acetylaspartate (NAA), taurine (Tau), glycerophosphocholine + phosphocholine (GPC+PCh), creatine + phosphocreatine (Cr+PCr), and Glu+Gln.

The statistical analysis Figure 8 showed increases in GABA, Gln, GSH, and Tau from Day 24 to 35, returning to baseline levels by Day 77 (Figure 8). In contrast, NAA exhibited the opposite pattern, being significantly reduced at Days 24–35 before recovering. Ins decreased at D24, rose at D35–D49, and then slightly declined by Day 77, while remaining elevated compared to baseline.

**Figure 8:**
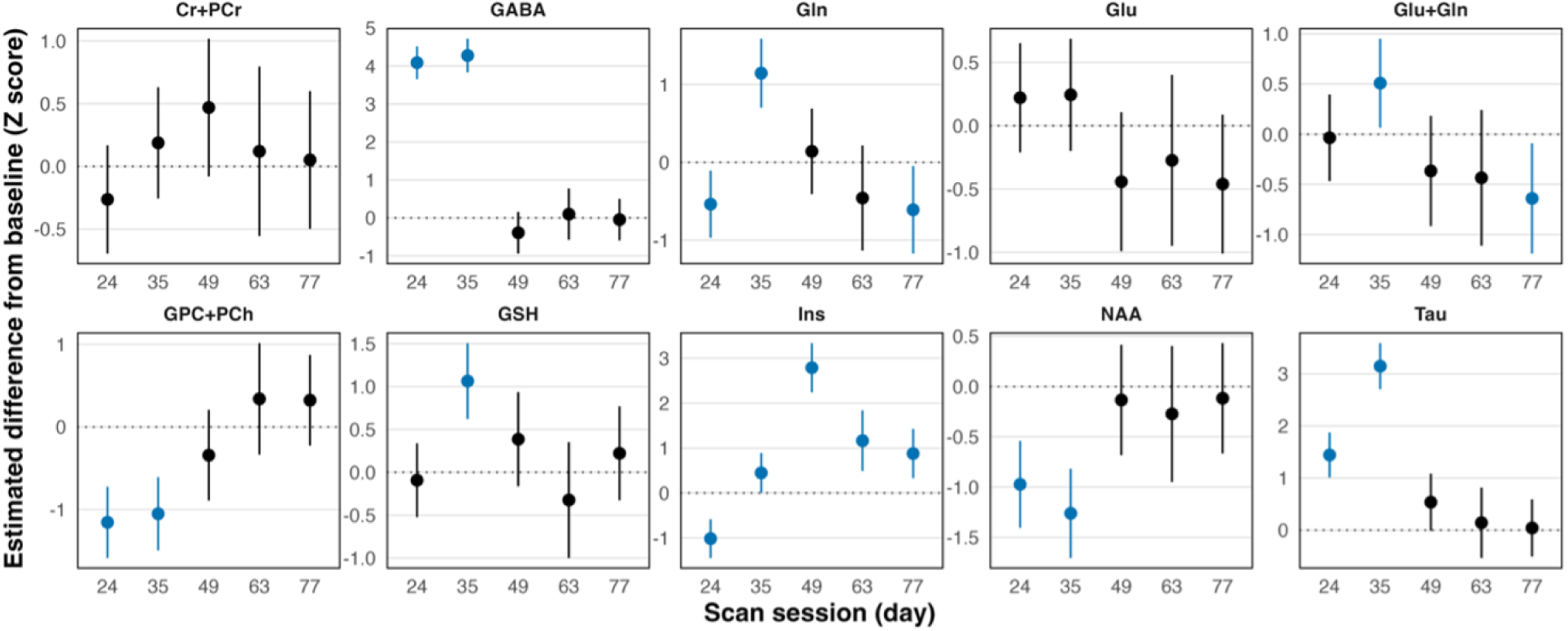
Differences from baseline in MRS metabolites within the corpus callosum. Values were normalised using the mean and standard deviation of the respective values for each metabolite from the baseline scans. A mixed-effects model was fit with a random intercept per subject, and metabolite and the interaction between metabolite and session as fixed effect predictors. The estimated marginal means (EMM) were calculated from each model fit, and 95% confidence intervals are shown. Šidák’s correction was applied to account for the 10 metabolite comparisons within a session. An EMM interval that did not include 0 was considered significantly different from baseline (shown in blue).

### Histology

A subset of mice from the in vivo study (see Table 2), together with non-cuprizone controls, underwent histology and immunohistochemistry at Days 24, 35, 42, and 77; representative images are shown in Figure 10. Quantification was performed in two corpus callosum ROIs and one cerebellar ROI (Figure 9).

**Figure 9:**
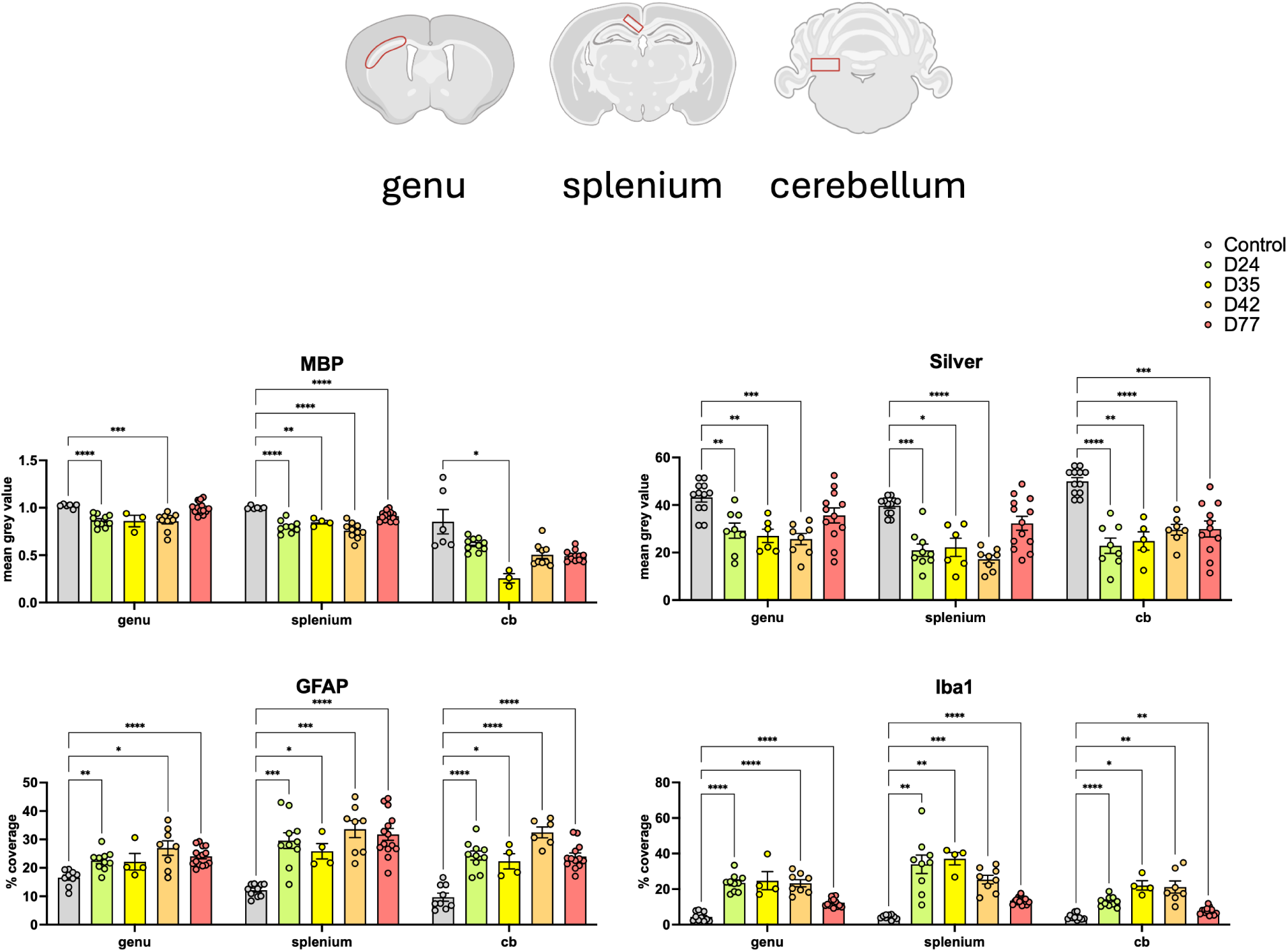
Histological characterisation of brain tissue from cuprizone-treated mice. Top panel: Regions-Of-Interest (ROIs) used for histological image analysis (red). Bottom panel: Analysis of myelin (myelin basic protein; MBP), nerve fibres (silver), astrocytes (GFAP), and microglia (Iba1) markers are shown from the corpus callosum lateral genu, medial splenium and cerebellum ROIs. Each data point represents the average of at least 4 images acquired from a given ROI from one mouse. Mixed ANOVA and Dunnett’s post hoc tests were used to compare each cuprizone group to control; ****p *<* 0.0001, ***p *<* 0.001, **p *<* 0.01, *p *<* 0.05.

**Figure 10:**
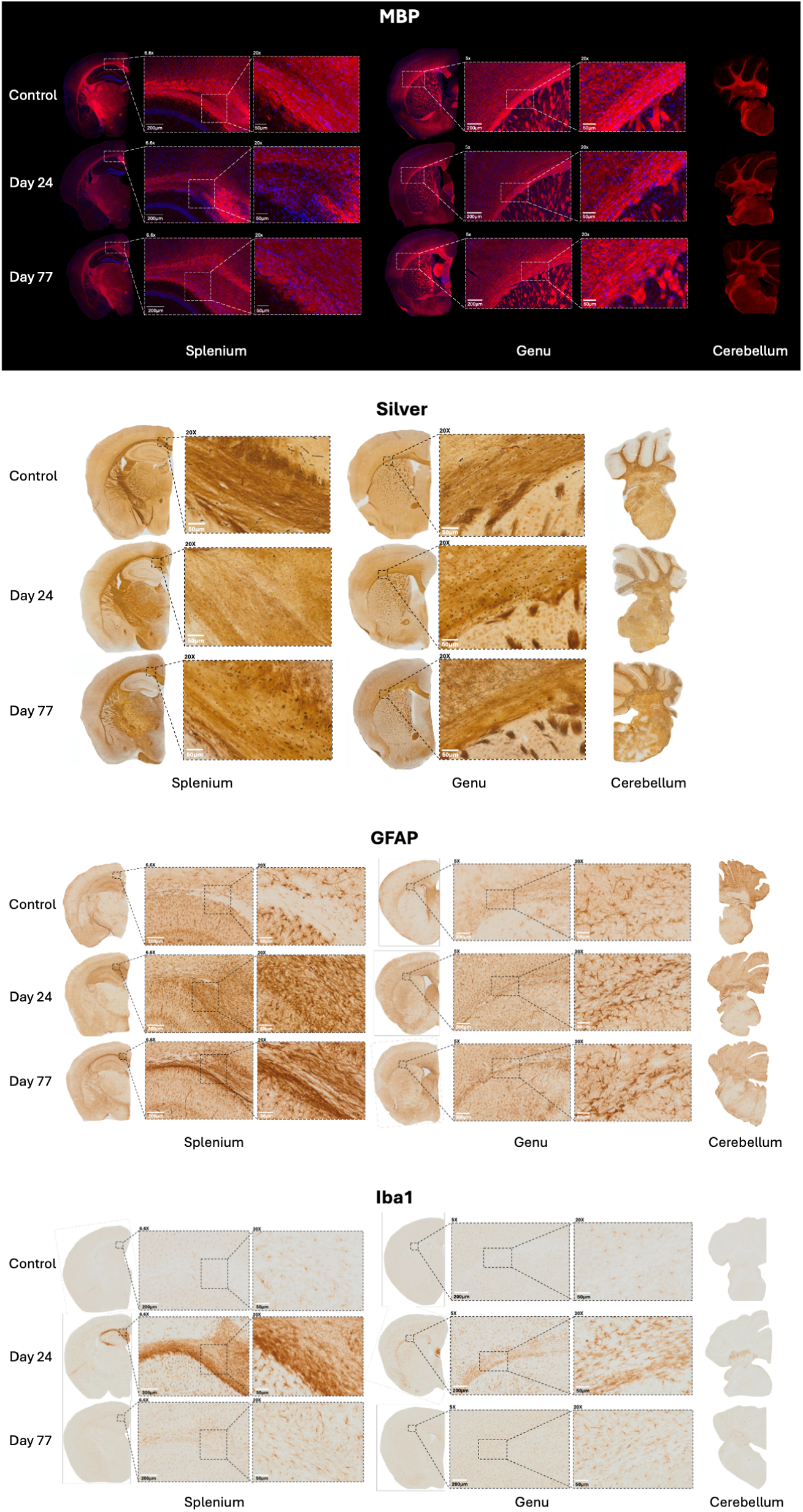
Representative histological images of brain tissue from cuprizone-treated mice(Day 24 and 77) and matched controls. Histological markers of myelin (myelin basic protein; MBP), nerve fibres (silver), astrocytes (GFAP), and microglia (Iba1) markers are shown in the corpus callosum medial splenium, lateral genu, and cerebellum regions of interest (ROIs).

Two myelination-related measures — MBP immunoreactivity and silver staining — showed decreases across all three ROIs that persisted through at least Day 42, with apparent recovery by Day 77 in the corpus callosum but not the cerebellum. Changes in silver staining were qualitatively stronger and more robust than those in MBP, although the spatial patterns were similar.

Inflammatory markers (GFAP for astrocytes and Iba1 for microglia) were significantly increased, consistent with reactive gliosis. GFAP increases were significant in all regions, except at the early time points (Days 24–35) in the genu, and remained elevated at Day 77. Iba1 signal increased prominently by Day 24 and then declined modestly, remaining significantly increased at Day 77 in the corpus callosum but not in the cerebellum. Notably, the temporal profiles diverged: Iba1 increased and resolved earlier, whereas GFAP rose more gradually and remained higher at Day 77.

## Discussion

In earlier work we characterised widespread brain changes in the mouse brains ex vivo, after 5 weeks of cuprizone exposure, identifying alterations in regional brain volume, DTI metrics, *T*_1_, *T*_2_, and myelin water fraction (Wood et al., 2016). Building on this, we now present a longitudinal in vivo study of the same inflammatory demyelination model, using updated and clinically translational MR-based metrics and a larger cohort. Mice were imaged longitudinally over 11 weeks, with cuprizone withdrawn at week 5 to capture both demyelination and subsequent remyelination, which is thought to nearly complete within 6 weeks after cessation of cuprizone (Matsushima and Morell, 2001; Kipp et al., 2009; Sachs et al., 2014; Tagge et al., 2016; Vega-Riquer et al., 2019). We employed quantitative, whole brain multi-modal MRI (MPM, DTI, volumetric imaging), and single voxel MRS at each timepoint, and we additionally acquired high resolution ex vivo imaging data at one timepoint (day 42). To improve sensitivity, in vivo data were pooled from three separate experiments. Our results corroborate previous findings while also revealing novel changes. Although histological characterisation was also performed, the key message is that MR-based characterisation is ideally suited to this model: it is non-invasive, faster, and—while less specific—often more sensitive than histology, which nevertheless remains the gold standard for cellular markers.

Quantitative MRI (MTsat*δ*, *R*_1_) revealed early, widespread demyelination beginning in the corpus callosum and cerebellar nuclei by Day 24 and extending to cortical and striatal regions by Day 35; abnormalities only partially resolved and remained evident through Day 77 (6 weeks after cuprizone cessation). Effects involved both grey and white matter, with MTsat*δ* changes qualitatively more extensive than *R*_1_. DTI demonstrated microstructural disruption, with a complex pattern of FA increases and decreases across grey and white matter, whereas MD changes were modest and largely transient. TBM showed persistent volumetric remodelling—enlargement of the hippocampus and cerebellar nuclei accompanied by cortical and cerebellar cortical volume loss—still present at Day 77. MRS in the corpus callosum revealed increased GABA and taurine and decreased NAA and choline-containing metabolites (glycerophosphocholine + phosphocholine) during cuprizone exposure (Days 24–35), most of which normalised by the end of treatment, while inositol remained elevated. Corroborative histology, on the other hand, indicated near-restoration of myelination by Day 77 (MBP and silver staining), but persistent reactive gliosis, with elevated astrocytic and microglial markers.

To further corroborate the in vivo imaging, we also examined ex vivo mouse data collected after 42 days of cuprizone treatment (7 days longer than the in vivo cohort). While the slightly different cuprizone treatment duration limits a direct comparison, the datasets are sufficiently related to justify a qualitative comparison. In the ex vivo samples, we acquired *R*_1_ and DTI, along with volumetric MRI (see below), but not MT. Here, both *R*_1_ and MD showed high sensitivity to widespread cuprizone-induced effects, consistent with our earlier ex vivo work (Wood et al., 2016) where *R*_1_ and MD changes appeared similar to each other in regions of pronounced pathology. The affected clusters in the present ex vivo dataset are larger than those reported in (Wood et al., 2016) likely due to improved scanner hardware, greater sample size, and the extended duration of cuprizone exposure.

Assuming the in vivo and ex vivo cohorts in this study developed comparable pathology (as supported by histology; manuscript in preparation), we noted an apparent discrepancy: DTI and *R*_1_ changes were more pronounced ex vivo than in vivo. While post-mortem biophysics (death, fixation, temperature) may contribute, a more likely explanation is reduced in vivo sensitivity due to motion, limited scan time, and lower resolution. For in vivo *R*_1_ mapping, the dual flip angle approach implemented in MPM is fast but susceptible to bias—especially at high field where B0 and *B*_1_+ inhomogeneities are greater—whereas the MP2RAGE sequence that was used for ex vivo *R*_1_ mapping is largely insensitive to *B*_1_+ inhomogeneities. Although we applied AFI-based *B*_1_+ correction, physiological motion and time constraints inherent to in vivo imaging lowered data quality relative to ex vivo *R*_1_. Likewise, our in vivo DTI used a fast EPI readout that is more artefact prone than the conventional spin-warp approach used for ex vivo DTI; ex vivo imaging also permitted longer acquisitions, higher spatial resolution, and improved SNR. More comparable *R*_1_ and DTI changes may have been observed in vivo and ex vivo had more comparable pulse sequences been used. Notably, the spatial distribution of *R*_1_ changes in the Day 42 ex vivo cohort (Figure 7) closely matched MTsat*δ* alterations in Day 35 and 49 in vivo (2, 6), suggesting these metrics detect the same pathology and, in this study, provided the most sensitive readouts for cuprizone-induced changes ex vivo and in vivo, respectively. This is relevant since studies sometimes interchangeably use in vivo and ex vivo approaches to characterise pathology. Although beyond the scope of this study, a more standardised, systematic evaluation of these imaging metrics with underlying biology would be valuable.

The changes we observed in MTsat*δ*, *R*_1_, and DTI metrics, resulting from cuprizone-induced pathology were remarkably brain-wide. Unlike many previous investigations that focused on predefined ROIs such as the corpus callosum, we imaged the whole brain and conducted the analyses voxelwise without regional restrictions. One of the central aims was to assess the extent and spatial pattern of pathology, rather than limiting the analysis to canonical regions. As expected, the corpus callosum was the most affected structure, but we also observed pronounced effects in the fimbria fornix and internal capsule, both major white matter tracts. As summarized in Figure 6 this approach revealed changes not only in white matter but also in grey matter regions. Several grey matter regions were impacted, including the cerebellar nuclei, cortex, thalamus, and striatum, each showing slightly different temporal profiles. These findings corroborate the emerging evidence that cuprizone affects the entire brain as has been seen for example by (Fjaer et al., 2013), who detected decreases in magnetization transfer ratio (MTR), a metric related to MTsat*δ*, in the cortical and subcortical grey matter in the mice. Although the authors did not look past 2 weeks after cuprizone, they also showed a high correlation between MTR and myelination. Several other studies (Petiet et al., 2016; Khodanovich et al., 2019; Guglielmetti et al., 2020; Hertanu et al., 2023) have explored changes in MRI signal in areas other than the corpus callosum although most do not report voxelwise maps of significant clusters as we do, limiting direct comparison.

In addition to the typical tissue microstructural measures (MTsat*δ*, DTI, *R*_1_) we also conducted whole brain tensor-based morphometry (TBM) analysis of our structural MRI to characterise regional volume changes compared to baseline in cuprizone-treated mice. TBM is a voxelwise method that quantifies local expansion or contraction from the deformation fields required to align individual images to a common template. This differs from voxel-based morphometry (VBM), which is more commonly applied in human studies and estimates tissue density at each voxel after segmentation into grey matter, white matter, or CSF (Ashburner and Friston, 2000). Because grey/white matter contrast is much lower in mice, segmentation is unreliable; TBM therefore provides a more robust and segmentation-free approach. In human MS, morphometry consistently revealed widespread brain atrophy linked to age and disease progression, correlating with cognitive decline and disability (Giorgio et al., 2008; Rothstein, 2020).

In our longitudinal in vivo data, TBM revealed a robust, widespread pattern of volume decreases across cerebral and cerebellar cortices that persisted even 6 weeks after cuprizone withdrawal. By contrast, regions strongly affected by both demyelination and inflammation, such as the corpus callosum and cerebellar nuclei, showed striking volume increases that were relatively stable over time, with modest attenuation at later stages. Subcortical regions including brainstem, thalamus, amygdala, and striatum displayed more complex trajectories, in some cases shifting from volume loss to gain. These findings are remarkable yet difficult to interpret without additional histological or molecular validation. Cortical atrophy may reflect grey matter demyelination, axonal degeneration, or synaptic/cellular loss (Giorgio et al., 2008; Bergsland et al., 2018; Ponticorvo et al., 2021), but the pattern does not align with other markers (MTsat*δ*, *R*_1_), suggesting additional processes are involved.

It will be important to determine whether these TBM changes reverse or persist once remyelination is complete, the latter potentially reflecting irreversible tissue loss. Addressing this question would require a longer study extending until remyelination is fully achieved and confirmed histologically, which was not feasible within the scope of the present work. Volume increases in the corpus callosum and cerebellar nuclei are most plausibly explained by inflammation-related processes, such as microglial or astrocytic proliferation, cellular swelling, and water accumulation (Uher et al., 2021; Wang et al., 2024). The dynamic changes in subcortical regions likely represent a combination of degeneration, cell loss, and inflammation. Future studies in models of pure demyelination or pure inflammation would be needed to disentangle these contributions.

Interestingly, we did not observe the same pattern in ex vivo TBM at day 42, as either Day 35 or 49 in vivo, although some similarities were apparent. Cortical decreases were largely absent ex vivo except in focal regions of cingulate, retrosplenial, and posterior cortex, while the cerebellum and thalamo-striatal areas showed partially overlapping effects. The most consistent findings across both in vivo and ex vivo were robust increases in cerebellar nuclei and hippocampus, which we again speculate reflect inflammatory changes. The absence of widespread cortical atrophy in ex vivo data may be due aforementioned biophysical tissue changes that result from death and fixation which induces shrinkage. If shrinkage disproportionately affected controls (with cuprizone brains potentially already somewhat atrophied), group differences may have been reduced.

Our prior ex vivo study showed a broadly similar pattern, albeit with smaller magnitude changes, likely reflecting differences in hardware, sample size, and processing (Wood et al., 2016).To our knowledge, this is the first application of whole-brain, longitudinal morphometry in the cuprizone mouse model, providing new insights into the regional dynamics of tissue loss and inflammation in this model.

We detected extensive temporal changes in MRS metabolites in cuprizone mice, with only 2 of the 10 measured metabolites or metabolite combinations remaining unchanged. One notable finding was an early increase in GABA (Days 24 and 35). To our knowledge, this has not previously been reported in the corpus callosum during cuprizone ingestion, potentially because GABA MRS is difficult to quantify without spectral editing (Rotaru, 2023). However, consistent with our results, elevated GABA has been reported within the first week of cuprizone exposure using other methods (Biancotti et al., 2008; Hayakawa et al., 2019). The significance of this increase remains uncertain, although GABA is known to regulate cell proliferation and immune function. Oligodendrocytes and OPCs both synthesise GABA and express GABA receptors, with receptor-specific effects: GABA_B_ receptor activation promotes OPC differentiation (Serrano-Regal et al., 2020; Bai et al., 2021), whereas GABA_A_ receptor activation can reduce OPC numbers, oligodendrocyte survival, and myelination in slice cultures (Hamilton et al., 2016). Thus, increased GABA may reflect an early endogenous repair response, potentially linked to the OPC proliferative phase that becomes prominent around the fourth week of cuprizone-induced demyelination (Brousse et al., 2015), while highlighting the need for GABA homeostasis in oligodendrocyte lineage function. In humans, MRS using the MEGAPoint Resolved Spectroscopy Sequence technique reported regional decreases in GABA in relapsing–remitting or progressive MS, correlating with worse cognitive performance (Cawley et al., 2015; Cao et al., 2018). In contrast, PET study reported increased GABA receptor density in deep grey matter that was associated with preserved cognitive function (Huiskamp et al., 2023). Enhanced GABAergic tone has also been linked to reduced active MS lesions during pregnancy (Kalakh and Mouihate, 2019), potentially via GABA_A_ receptor–mediated immunomodulation (Tian and Kaufman, 2023). Overall, our findings support GABA as an early biomarker of cuprizone-induced damage, but further work is needed to clarify its functional significance.

Other neurotransmitter-related metabolites of interest in this model include glutamate and glutamine. In contrast to previous reports (Orije et al., 2015; Genovese et al., 2021), we did not detect significant changes in glutamate despite the high sensitivity of our MRS acquisition. Instead, we observed an increase in Glx (glutamate + glutamine) at Day 35, driven primarily by glutamine fluctuations (decreased at Day 24, increased at Day 35). As glutamine is the precursor of glutamate, this pattern may indicate altered glutamatergic cycling. The elevation in glutamine/Glx at Day 35 coincides with peak pathology and could reflect excitotoxic mechanisms, while the subsequent decrease in glutamine by Day 77 suggests a dynamic trajectory of glutamatergic cycling as pathology progresses.

Of other metabolites, choline was reduced early (Days 24-35) but later normalized, consistent with transient membrane breakdown during acute demyelination, as previously reported (Orije et al., 2015; Xuan et al., 2015). Taurine was also increased at early stages, in agreement with prior reports (Orije et al., 2015; Genovese et al., 2021), though this response may be more pronounced in rodents than humans (Sturman, 1986; Puka et al., 1991). NAA levels decreased early (Days 24-35) and recovered later, a common finding in cuprizone studies (Sachs et al., 2014; Orije et al., 2015; Genovese et al., 2021). Finally, inositol showed the most pronounced changes, with an initial decrease (Day 24) followed by sustained increases (Day 35 onwards). This biphasic response likely reflects a shift from acute astrocytic/metabolic dysfunction to chronic astrogliosis, glial scarring, and altered osmotic regulation. Previous in vivo work on inositol in cuprizone is scarce, with only (Sachs et al., 2014) reporting a decrease at 6 weeks. Our data extend these observations by demonstrating a biphasic response, although further work could explicitly model this trajectory.

Our histological analysis broadly corroborated the imaging: we observed marked reductions in MBP and silver staining (reflecting demyelination) during cuprizone exposure, with restoration evident by Day 77 in the corpus callosum but not the cerebellum. This diverged from persistent MTsat*δ* and *R*_1_ abnormalities in vivo, suggesting that MRI may detect residual microstructural alterations below the sensitivity or sampling of our histology. Corroborating this interpretation, a study combining ultrashort-TE with MT (UTE-MT), demonstrating strong histological correlation with myelination, reported incomplete remyelination by 6 weeks post-cuprizone, echoing our findings (Guglielmetti et al., 2020). Moreover, a behavioural study established that cuprizone-lesioned mice appear to have long lasting impairments in their sensory and motor function, remaining at least 6 weeks after cuprizone, which could match our finding of prolonged pathology (Tomas-Roig et al., 2019). On the other hand, it is known that magnetization transfer ratio imaging (related to MTsat*δ* metric) can be influenced by processes beyond demyelination, such as axonal pathology and inflammation (Tagge et al., 2016). Given that axonopathy has not been widely reported in this model (Merkler et al., 2005), a major contribution from axonal injury seems unlikely, so we did not test for it in this study. However, given that gliosis is a prominent feature of this model, an inflammatory contribution to the MTsat*δ* signal cannot be excluded.

Robust reactive gliosis was also detected by histology—both microglia and astrocyte signals were elevated and remained so at Day 77—aligning with sustained inositol increases on MRS and volume enlargements in the corpus callosum and cerebellar nuclei on TBM, consistent with inflammation-related processes. Notably, inflammation outlasted (re)myelination: while myelin-sensitive histological metrics partially normalised, gliotic markers persisted, in keeping with reports that remyelination can approach completion within weeks of cuprizone withdrawal whereas gliosis resolves more slowly (Gudi et al., 2014; Praet et al., 2014). Taken together, the spatial concordance between ex vivo *R*_1_ (Day 42) and in vivo MTsat*δ* (Days 35–49) supports the view that MTsat*δ* and *R*_1_ index demyelination–remyelination and, on MRI, remyelination appears incomplete by Day 77. On the other hand, MRS (inositol) and TBM appear to provide complementary sensitivity to inflammatory remodelling.

Nevertheless, the relative contribution of inflammation and demyelination in our imaging data is unclear, as is the contribution of the secondary pathology – cell death, axonal derangement, plastic rewiring of the brain in response to widespread pathology. Indeed, it has not yet been confirmed that all regions where we detect MRI changes are substantially demyelinated, since neither we nor others conducted whole brain histology. It is a disadvantage of histology that more subtle changes may go undetected and that studies are often underpowered to detect smaller differences because animals must be killed to conduct histology. Further methodological downsides of histology include relying on manual tissue processing, which is prone to artifacts, tissue shrinkage, and usually being limited to one or two markers. By contrast, MRI provides non-invasive, multi-modal readouts, capturing several indirect markers simultaneously without altering the tissue. While MRI is less specific than histology, it may be more sensitive to subtle microstructural changes that are challenging to detect histologically. Importantly, this is also the strength of imaging: MRI can guide subsequent post-mortem studies, whether in the same animals (though at the expense of longitudinal imaging) or in parallel cohorts.

This study has several limitations. First, histology was limited in scope and not systematically compared to nor co-registered with MRI for voxelwise correlation, limiting direct biological interpretation. Second, we lacked separate time-matched control cohorts and instead compared scans at timepoints to baseline; although large effects were detected compared to baseline, slow drifts in putative “control” signal cannot be fully excluded (Tagge et al., 2016). We did observe whole-brain volume increases in the mice over time (Figure S5), so some imaging changes could reflect age-related growth; however, robust, anatomically plausible effects were already present at Days 24–35—before substantial global volume increases—and later regional patterns largely followed those of earlier timepoints, evidence against major age-related confounds. In accordance with the 3Rs, we opted to limit animal use and therefore did not include an additional non-cuprizone control group. Third, the cohort was predominantly male, with too few females for a powered sex analysis; while demyelination markers did not appear to differ markedly by sex in our dataset, as has been reported by others (Taylor et al., 2010), formal sex-by-metric testing across all modalities we used would be important. Finally, in vivo acquisitions faced motion and scan-time constraints that influenced our choice of sequences, and ex vivo data were acquired at a single time point without MT and after a slightly longer cuprizone exposure—all these factors complicate direct modality-to-modality comparisons.

## Conclusion

We provide a comprehensive, longitudinal, in vivo characterisation of cuprizone-induced demyelination and partial repair, complemented by ex vivo imaging and histology. Optimised multimodal MRI (MTsat*δ*, *R*_1_, DTI, TBM) and corpus callosum MRS sensitively tracked early myelin loss, evolving microstructural and volumetric changes, and a distinct metabolic signature, suggesting that myelin recovery remains incomplete six weeks after cuprizone and that glial-driven inflammation can outlast myelin repair. Rather than exhaustively dissecting molecular mechanisms, our aim was to establish a robust in vivo assay for testing experimental therapeutics: we found MTsat*δ* (in vivo) and *R*_1_ (ex vivo) were the most sensitive readouts of cuprizone pathology, with MRS/TBM providing complementary information on inflammatory and structural remodelling.

## Abbreviations

**Abbreviation Definition**
AD: axial diffusivity
ANOVA: analysis of variance
ANTs: advanced normalisation tools
B1: radiofrequency magnetic field
Cb: cerebellum
CC: corpus callosum
CCFv3: common coordinate framework version 3
COMPOSER: combining phase data using a short echo-time reference scan
CPZ: cuprizone
Cr+PCr: creatine + phosphocreatine
DAB: 3,3-diaminobenzidine
DTI: diffusion tensor imaging
DWI: diffusion-weighted imaging
ECSTATICS: ESTimating the Apparent Transverse relaxation time from Images with different ContrastS
EMM: estimated marginal means
FA: fractional anisotropy
FWE: family-wise error
FID-A: free induction decay analysis appliance
FLAIR: fluid-attenuated inversion recovery
FOV: field of view
FSL: FMRIB software library
GABA: *γ*-aminobutyric acid
GFAP: glial fibrillary acidic protein
Gln: glutamine
Glu: glutamate
Glu+Gln / Glx: glutamate + glutamine
GPC+PCh: glycerophosphocholine + phosphocholine
GSH: glutathione
H_2_O_2_: hydrogen peroxide
Iba1: ionized calcium binding adaptor molecule 1
IHC: immunohistochemistry
IMS: industrial methylated spirits
Ins: inositol
LC: linear combination
MBP: myelin basic protein
MD: mean diffusivity
MP2RAGE: magnetisation prepared 2 rapid acquisition gradient echoes
MPM: multiparametric mapping
MRI: magnetic resonance imaging
MRS: magnetic resonance spectroscopy
MS: multiple sclerosis
MTsat: magnetisation transfer saturation
MTsat*δ*: change in MTsat
MTw: magnetisation transfer-weighted
NAA: N-acetylaspartate
NAAG: N-acetylaspartylglutamate
OPCs: oligodendrocyte precursor cells
PD: proton density
PD*: effective proton density
PDw: proton density-weighted
PRESS: pointed resolved spectroscopy
QUIT: quantitative imaging tools
*R*_1_: longitudinal relaxation rate
R2*: effective transverse relaxation rate
RARE: rapid acquisition with relaxation enhancement
RD: radial diffusivity
ROI: region of interest
RT: room temperature
SE-EPI: spin-echo echo planar imaging
SMP: skimmed milk powder
*T*_1_: longitudinal relaxation time
*T*_1_w: T1-weighted
Tau: taurine
TB1AFI: transmit B1 field map actual flip angle imaging
TBM: tensor-based morphometry
TBS: tris-buffered saline
TFCE: threshold-free cluster enhancement
UTE: ultrashort echo time
VAPOR: variable pulse power and optimized relaxation delays
VBM: voxel-based morphometry

## Supplementary material

**Figure S1:**
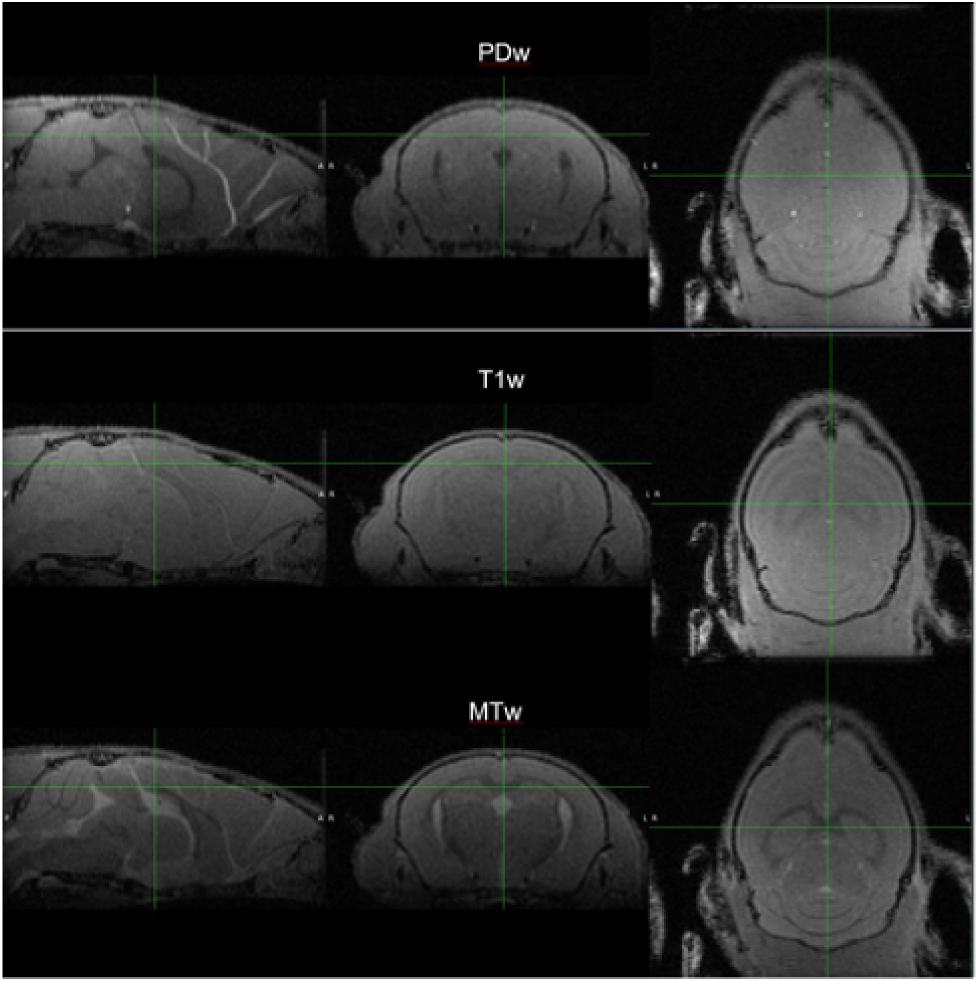
Representative multiparametric mapping (MPM) images from one mouse. Upper panel: proton density weighted (PDw) images; middle panel: *T*_1_ weighted images (*T*_1_w); lower panel: magnetization transfer weighted (MTw) images.

**Figure S2:**
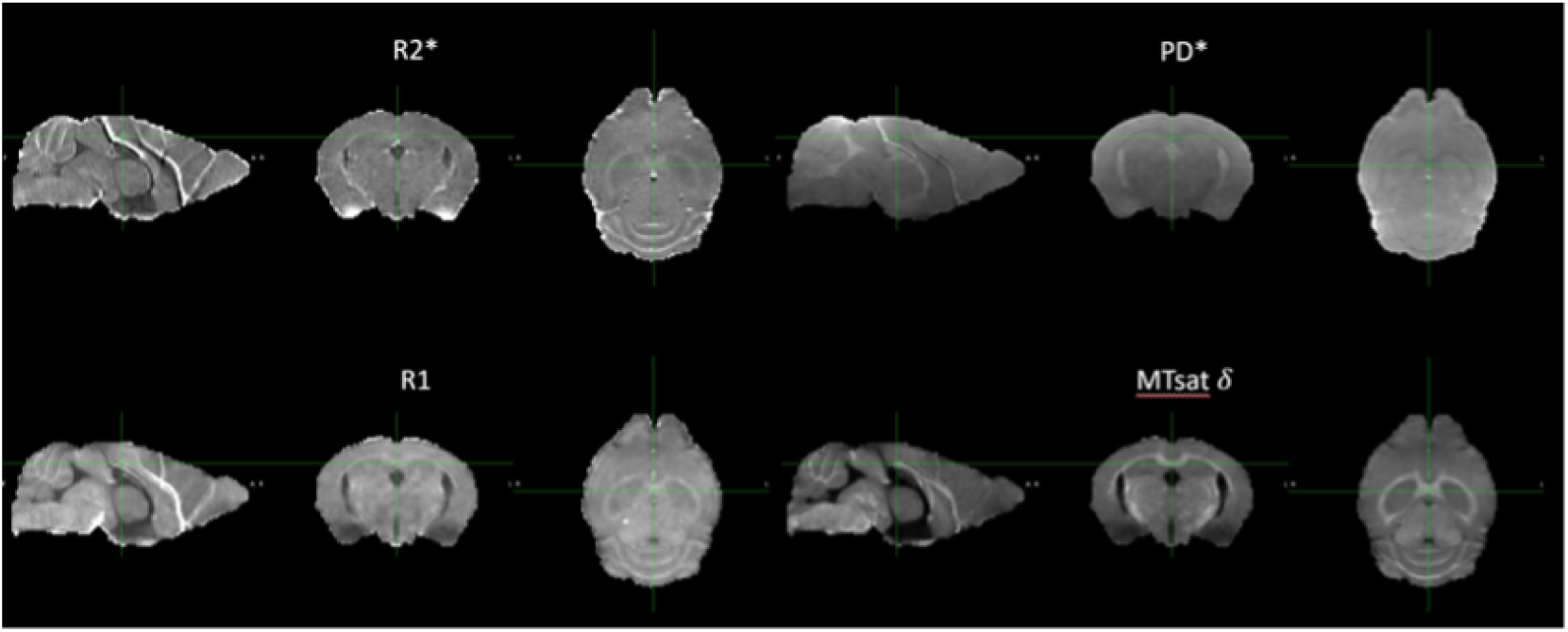
Representative multiparametric mapping (MPM) R2*, *R*_1_, PD* and MTsat*δ* maps from one mouse.

**Figure S3:**
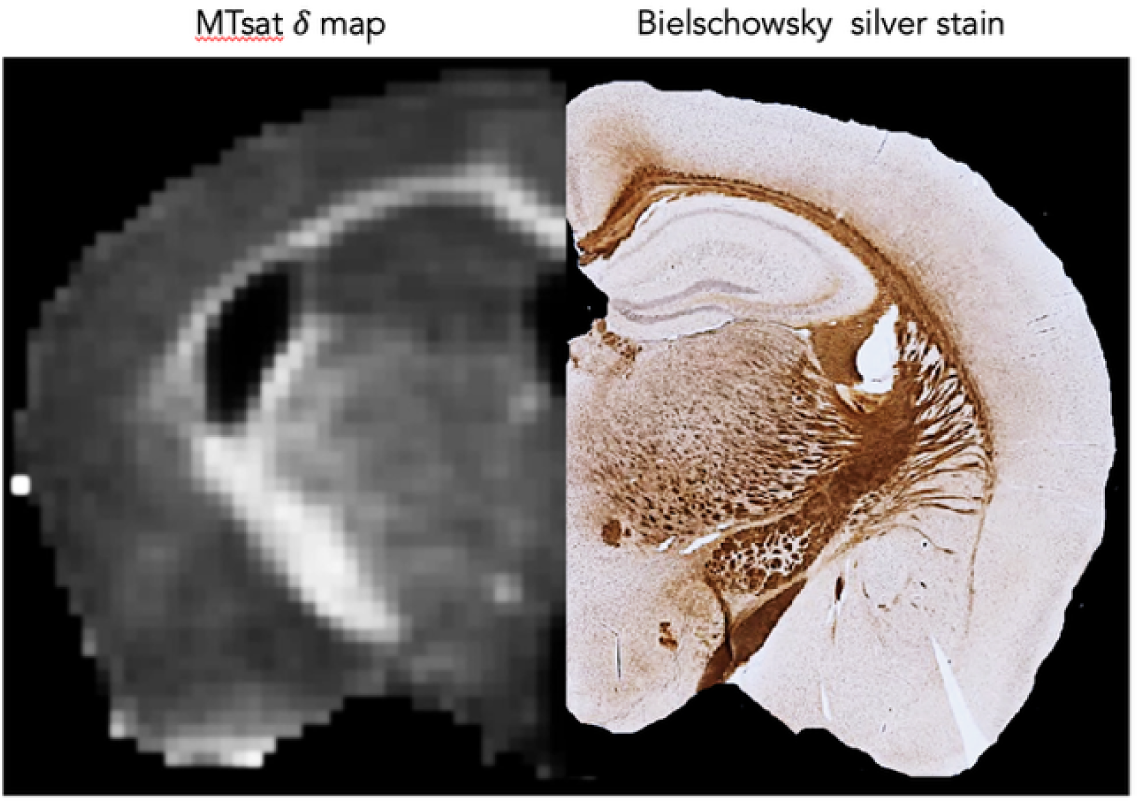
MTsat*δ* parameter approximates myelin Representative MTsat*δ* map from a mouse brain (left), showing hyperintense signals in myelinated regions, compared with a Bielschowsky silver stain highlighting axonal fibers (brown). A good qualitative correspondence is observed between the two images.

**Figure S4:**
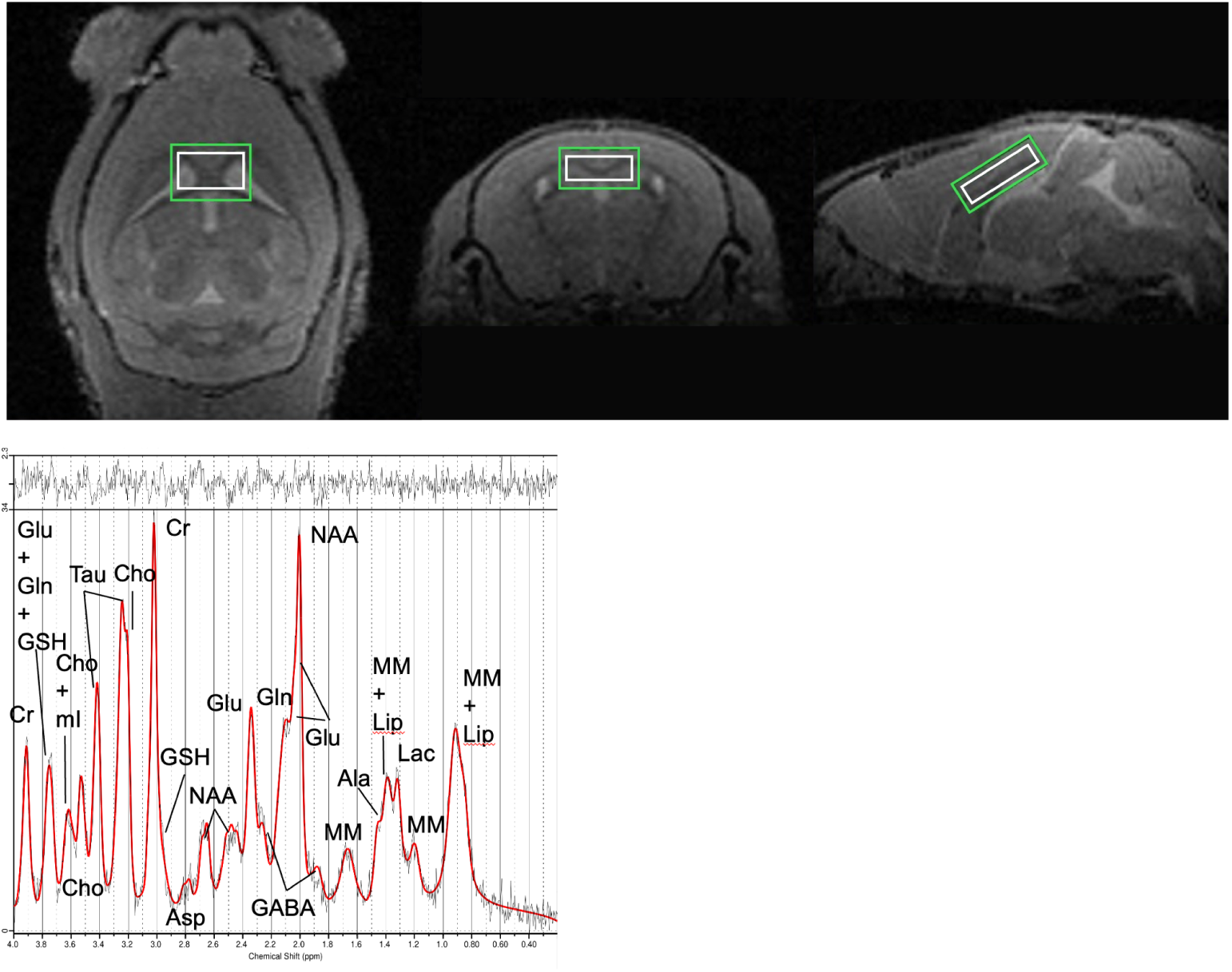
MRS voxel position and representative spectrum. Upper panel; voxel position for magnetic resonance spectroscopy (MRS). Voxel (white) and shim volume (green) position used for MRS scan. Lower panel; representative MR spectrum from one mouse (black = raw data; red = fitted data) with main metabolites labelled.

**Figure S5:**
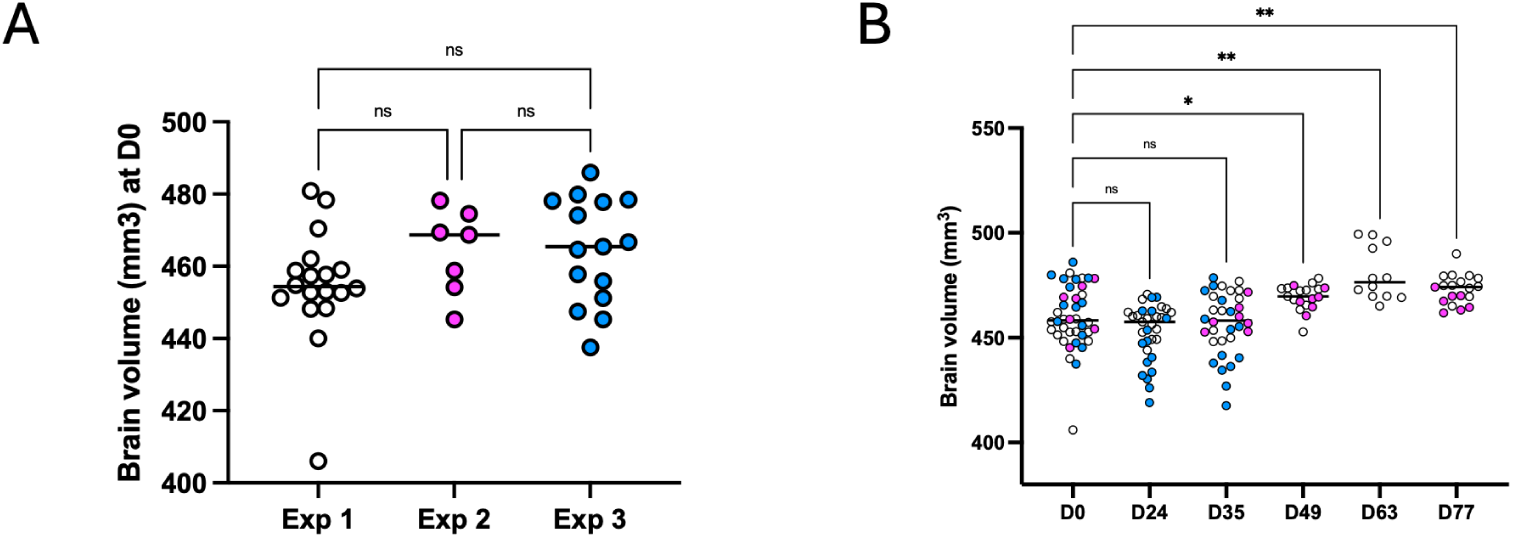
Whole brain volumes of all mice imaged in vivo. Whole-brain volumes for all mice included in the analysis, pooled across three independent experiments conducted with identical imaging parameters; at D0 baseline (A) and at all timepoints (B). Analysis was by (A) one way ANOVA and post-hoc Tukey between all groups, and (B) mixed ANOVA with post-hoc comparisons (Dunnett’s) versus D0 baseline; *p *<* 0.05, **p *<* 0.01. Each circle is a mouse with a horizontal line at the group median.

